# Brain-Wide Subnetworks within and between Naturally Socializing Typical and Autism Model Mice

**DOI:** 10.1101/2025.10.16.682530

**Authors:** Odeya Marmor, Renana Terner, Vivian Khuory, Shelly Ginzburg, Haitham Amal, Ariel Gilad

## Abstract

Social interaction is inherently asymmetric, requiring coordinated activity between non-homologous brain regions across individuals. However, the brain-wide dynamics underlying such inter-brain coordination remain poorly understood. We used multi-fiber photometry to simultaneously record from 24 brain regions in pairs of freely interacting mice, including a model of autism. Social interactions evoked widespread, dynamic activity across brains, with inter-brain synchrony, especially between non-homologous areas, exceeding intra-brain synchrony, particularly in dominant mice. Network analysis revealed three subnetworks: (1) Emotional, intra-brain enhanced in subordinates; (2) Sensory, spanning both mice; (3) Decision/consolidation, linking dominant prefrontal cortex to subordinate hippocampus. These subnetworks encoded dominance, identity, and interaction roles, and followed a clear temporal sequence around social events. In an autism model, socially evoked activity was hyperactive displaying mostly within brain synchrony but lacked inter-brain synchrony. Our results uncover dynamic inter-brain circuits as a hallmark of social behavior and reveal their disruption in autism.

## Introduction

Social behavior is among the most complex and evolutionarily conserved forms of interaction in mammals. Rather than being localized to isolated neural circuits, social information is increasingly recognized as emerging from coordinated activity across distributed, brain-wide networks. Human neuroimaging studies adopting whole-brain approaches have identified key regions involved in social processing, including the prefrontal cortex, amygdala, and others(*1*). However, these studies are limited by relatively low spatiotemporal resolution and constraints imposed by the scanner environment.

To more fully understand the complex neurobiology of social behavior, it is essential to study brain-wide activity during naturalistic, unrestrained interactions and, crucially, to measure neural dynamics across both interacting brains (but see(*2–4*) for studies using EEG from two interacting Humans). Animal models such as mice and bats offer an ideal platform for this, enabling high-resolution recordings during natural social behavior. While previous animal studies have provided important insights by focusing on specific brain regions such as the prefrontal cortex(*5–9*) nucleus accumbens(*10–12*), hippocampus(*13*, *14*) and amygdala (*15–17*), they largely center on isolated nodes or dyadic pathways e.g., thalamo-prefrontal or fronto-striatal circuits(*18–23*), leaving brain-wide interactions underexplored.

Recent technical advances have enabled simultaneous recordings from multiple brain areas during social contact, revealing the importance of distributed, large-scale networks(*24–26*). However, studies that map brain-wide dynamics across two interacting individuals remain scarce. This is a critical gap: social interactions are inherently asymmetrical, often involving distinct roles such as initiator and responder. Therefore, interactions may not be confined to homologous regions in each brain, but may instead involve cross-brain, cross-region communication, a concept not well captured in prior studies focusing on inter-brain interactions between homologous sites e.g., dual prefrontal cortex recordings(*5*, *8*, *9*, *13*, *27*).

An additional important aspect is how brain-wide dynamics are altered in autism spectrum disorder (ASD), where social interactions are critically impaired (*28–31*). Rather than being confined to specific loci, abnormalities in the autistic brain are increasingly recognized as distributed across brain-wide networks (*32–35*). Here we use a mouse model for ASD which has a genetic mutation in SHANK3 a scaffold protein in the postsynaptic density complex (*36*, *37*). Mutations in SHANK3 are strongly associated with ASD patients (*38*) and mutant mice (termed here as ASD mice) have been shown to display impaired sociability and other behavioral deficits (*39–41*). Human neuroimaging and rodent models alike reveal widespread alterations in functional connectivity, including excessive local coherence and disrupted long-range coupling across cortical and subcortical systems, e.g., amygdala, nucleus accumbens and prefrontal cortex(*32–34*, *42–44*). It is still debated whether ASD individual display hyper- or hypo-responses in different brain areas (*45–50*). How such network-level changes unfold during natural social interactions remains poorly understood, especially between two interacting partners. Investigating brain-wide and inter-brain dynamics during natural social interaction in models of autism therefore provides a critical opportunity to link distributed network abnormalities to atypical social behavior.

To address these limitations, we used multi-fiber photometry(*51*) to simultaneously record calcium dynamics from 24 brain regions from pairs of freely interacting normal and ASD mice. By recording across all pairs within groups of 4–5 mice, we captured brain-wide activity within and between individual brains during spontaneous social interactions. Normal mice display distinct functional subnetworks within and between brains, that emerge seconds before social contact, with patterns that differ based on individual characteristics such as social dominance. ASD mice display dysfunctional brain-wide networks with decreased inter-brain synchrony and enhance within-brain synchrony. These findings highlight a dynamic, asymmetric organization of brain-wide activity that underlies natural social behavior, providing a systems-level view of the social brain in action.

## Results

### Naturally socializing mice reveals complex social traits

To study brain-wide dynamics during natural social interaction we implemented multi-fiber photometry in pairs of freely moving mice within a behavioral arena (Fig. 1a). In total, we imaged all possible pairs within a group of mice (4 or 5 housed together; 6 or 10 pairs per group, 23 mice, 42 pairs in total; 12±1 10-minute sessions per mouse. 992 sessions in total). Both mice were tracked offline using DeepLabCut(*52*) and contact epochs were defined based on the Euclidean distance threshold (<8 cm for ≥0.5 s; Fig. 1b; See Methods). Each session had varying social dynamics depending on the relations within each pair of mice (Fig. 1c). On average, mice socially interacted 21±6% (mean±SEM) and rather similarly across all groups (Fig. 1d), with 24±11 social interactions per session, with a duration of 4.6±2.2 seconds per interaction. Next, we wanted to rank each mouse within its own group (n=4 or 5) based on the observed social dynamics. For each detected social interaction, we identified the initiating mouse based on approach velocity and tail-nose metrics (see Methods). The social rank of each mouse was defined as the ratio between initiation and being approached (averaged across all other mice in the group; Fig. 1e, f). Other behavioral parameters, including chase frequency, traveling velocity, cage occupancy and anxiety-related performance were also correlated to our social rank (Fig. S1). This dataset captures the rich and multidimensional dynamics of naturally socializing mice.

**Figure 1.**
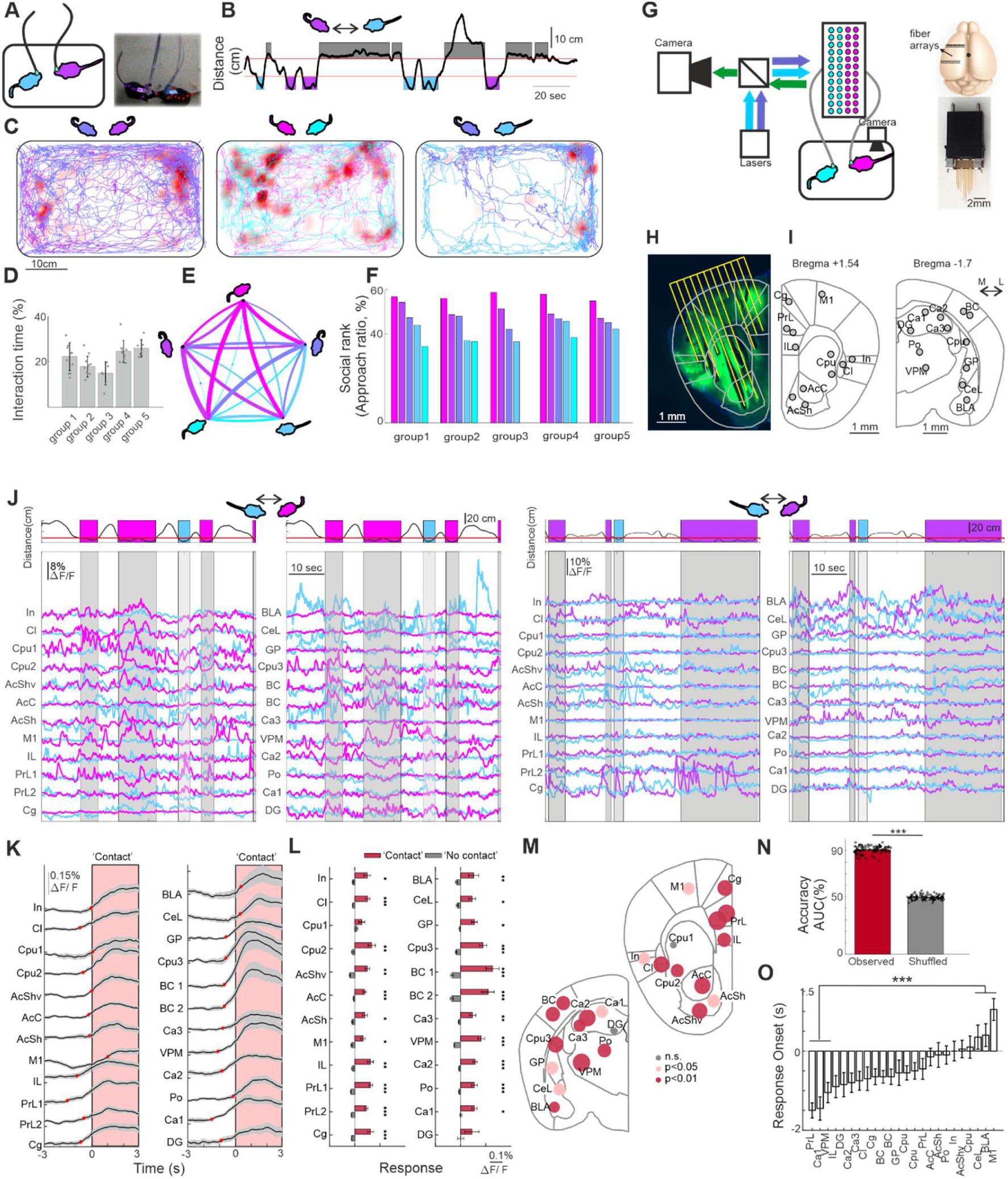
Brain-wide dynamics during natural social interaction. **a.** Schematic illustration of behavioral paradigm**. b.** Example of distance between mice during natural social interactions. Red lines indicate threshold for social interaction (’contact’; <8cm) and ‘no-contact’ (>15cm). Color of interaction depicts the initiating mouse. **c.** Example positions of two mice within the cage. Red shadow indicates social interactions. **d.** Social interaction time (%) for each group. error bars depict mean±SEM across recording days. Each dot represents one mouse. **e.** A diagram showing the ratio of initiating a social contact between each two mice within an example group. Thickness of each line indicates percentage of initiation. **f.** Relative rank of each group based on the initiation ratio derived from e, i.e., the ratio between initiating and being approached for each pair averaged across pairs for each mouse. Colors denote relative rank. **g.** Schematic illustration of multi-fiber setup and fiber positions from a top view. **h.** Coronal slices showing fiber tracks and green fluorescence of the calcium indicator (GCaMP6f). **i.** Fiber tip locations superimposed on two coronal sections, a frontal and a posterior one. The targeted areas are: Cg-cingulate cortex 1;PrL- prelimbic cortex, IL- infralimbic, M1-primary motor cortex, AcSh- Nucleus accumbens Shell, AcC- Nucleus accumbens core, Cpu- Caudate putamen, Cl- Claustrum, In- Disgranular insular cortex, DG- Hippocampus dentate gyrus, Ca1, Ca2, Ca3 - Hippocampus Cornu Ammonis 1, 2 and 3, Po- Posterior thalamic nuclear group VPM- Ventro posterior medial nucleus of thalamus, GP- the external boundary of gllobus pallidus, CeL- Centrolateral amygdala, BLA- Basolateral amygdala. **j.** Two examples showing neuronal responses (ΔF/F) in 24 areas as two mice freely move within the behavioral cage. Distance between mice displayed at the top, and social interactions (gray shaded areas) are colored based on the initiating mouse. **k.** Responses (ΔF/F) in 24 brain areas aligned on social contact (time 0). Error bars (shaded) depict mean±SEM across mice (n=23). Red dots indicate response onset for each brain area. **l.** Grand average responses averaged during social interaction (i.e., contact; red) and outside social interaction (i.e., no-contact; gray). Error bras as in k. *** *p <*0.001,\*\**p*<0.01,\**p*<0.05, Wilcoxon signed test with Bonferroni correction (n=23). **m.** Circle representation depicting the difference in responses between contact and no-contact periods divided by the sum SEM of both. The larger the circle the bigger the distance between contact and no-contact periods. Significance levels are color coded. **n.** The accuracy of an SVM classifier predicting contact or no-contact periods based on responses in 24 brain areas (Red; see Methods). Accuracy of the SVM on trial-shuffled data is depicted in the gray bar. Each gray dot represents one mouse. **\*\***\*p<0.001, Signed rank test. **o.** Response onset (in relation to social contact) in each brain area (sorted). Error bars as in k. ***p<0.001 Wilcoxon signed test.

### Brain-wide dynamics from two mice during natural social interaction

As described above, we used multi-fiber photometry to record brain-wide calcium dynamics (GCaMP6f) from both mice during natural social interaction. This approach enables simultaneous recording from 24 fiber sites across cortical and subcortical regions, including amygdala, prefrontal cortex, hippocampus and more (Fig. 1g-l; see Methods). Briefly, blue (473 nm) and purple (405 nm; control) light are interleaved, expanded and focused onto the fiber array while emitted green fluorescence from each fiber tip was captured by a CMOS sensor (Fig. 1g; see Methods). The fiber placements targeted both socially and non-socially related regions (Fig. 1I). At the end of experiments, brains were extracted, and fiber tip locations were histologically validated (Fig. 1h and Fig. S2; See Methods).

Fluorescence signals were normalized (ΔF/F) and corrected for hemodynamics artifacts (see Methods). Example traces from two mice during natural social interactions revealed rich brain-wide dynamics during social contacts (Fig. 1j). To quantify socially evoked response, we aligned the neuronal response in each brain area to social contact. Averaged across all mice, neuronal activity increased around contact events in nearly all regions, with varying onsets (Fig. 1k). Most regions showed significantly elevated activity during social contact epochs (mice< 8cm) compared to non-contact periods (> 15 cm; Fig. 1l, m; *p*<0.05; Wilcoxon signed-rank test; n=23 mice; Bonferroni corrected). These widespread responses extended beyond classically social areas to thalamus, Barrel cortex, hippocampus and striatum. Furthermore, a trained classifier (support vector machine) predicted social contact with high accuracy (90%±0.6%; mean±SEM) outperforming label-shuffled controls (Fig. 1n; see Methods). Next, we quantified response onset (in relation to social contact) as the first time frame crossing mean±3SD of baseline response calculated for each mouse and averaged across mice. Response onset varied across brain areas, with prelimbic and hippocampal areas displaying anticipatory responses (i.e., onset before social contact), significantly earlier than other brain areas such as amygdala (Fig. 1o, Wilcoxon signed-rank test, n=23 mice).

Because social contact coincides with movement, general exploration and sensory cues, we tested potential confounds: (1) To control movement, during non-contact periods, we compared slow (<1 cm/s) and fast (>4 cm/s) locomotion and found minor and mostly insignificant differences across the brain (Fig. S3. a–d). (2) To control general (non-social) exploration, we investigated brain-wide responses when mice explored stationary object and found no significant differences across the brain between object contact and no contact (Fig. S3. e). (3) In a separate cohort we imaged two mice freely moving in separate adjacent cages and pseudo-contact was defined by proximity, and no significant brain activity differences were observed (Fig. S3. f). Together, these results demonstrate that many brain regions respond during social contact, with diverse timing profiles, motivating to further dissect socially related subnetworks within and between two partners.

### Brain-wide subnetworks within and between dominant and subordinate mice

These experiments allow us to investigate the relationship between multiple brain areas between and within two mice. Thus, we first computed the full correlation matrix between all brain areas (within and between mice) during social contact and during ‘no-contact’ epochs (Fig. S4. a). A differential correlation matrix (Δr) was then calculated by subtracting the no-contact matrix from the contact matrix, per mice pair, and averaged across all pairs (n=42) while preserving social hierarchy order (dominant always on the left; Fig. 2a). Each matrix element represents the change in correlation (Δr) between two brain regions during contact relative to no-contact. Positive Δr values reflect enhanced synchrony during social contact; negative values reflect the opposite (see Methods). In general, we observed significantly positive values between brains (averaging the top right quadrant in Fig. 2a), indicating increased inter-brain correlations during social contact (Fig. 2b; *p*<0.01; Signed rank test). In contrast, within-brain Δr values (for both dominant and subordinate mice) were not significantly different from zero (p>0.05; Wilcoxon signed-rank test, n=42) with a trend for higher values in the subordinate mouse as compared to the dominant mouse (p=0.054). To test whether increased inter-brain synchrony could arise from shared time-locked external inputs, we performed a control analysis by shuffling contact epochs across unpaired mouse brains (i.e., aligning contact activity in one mouse with that of a different, non-interacting mouse; see Methods; (*8*)). The inter-brain Δr values in the unpaired control were not significantly different than 0 and were significantly lower than the observed Δr values (*p*<0.05; Wilcoxon signed-rank test; Fig. 2b). Thus, social contact evokes widespread synchrony between the two interacting brains.

**Figure 2.**
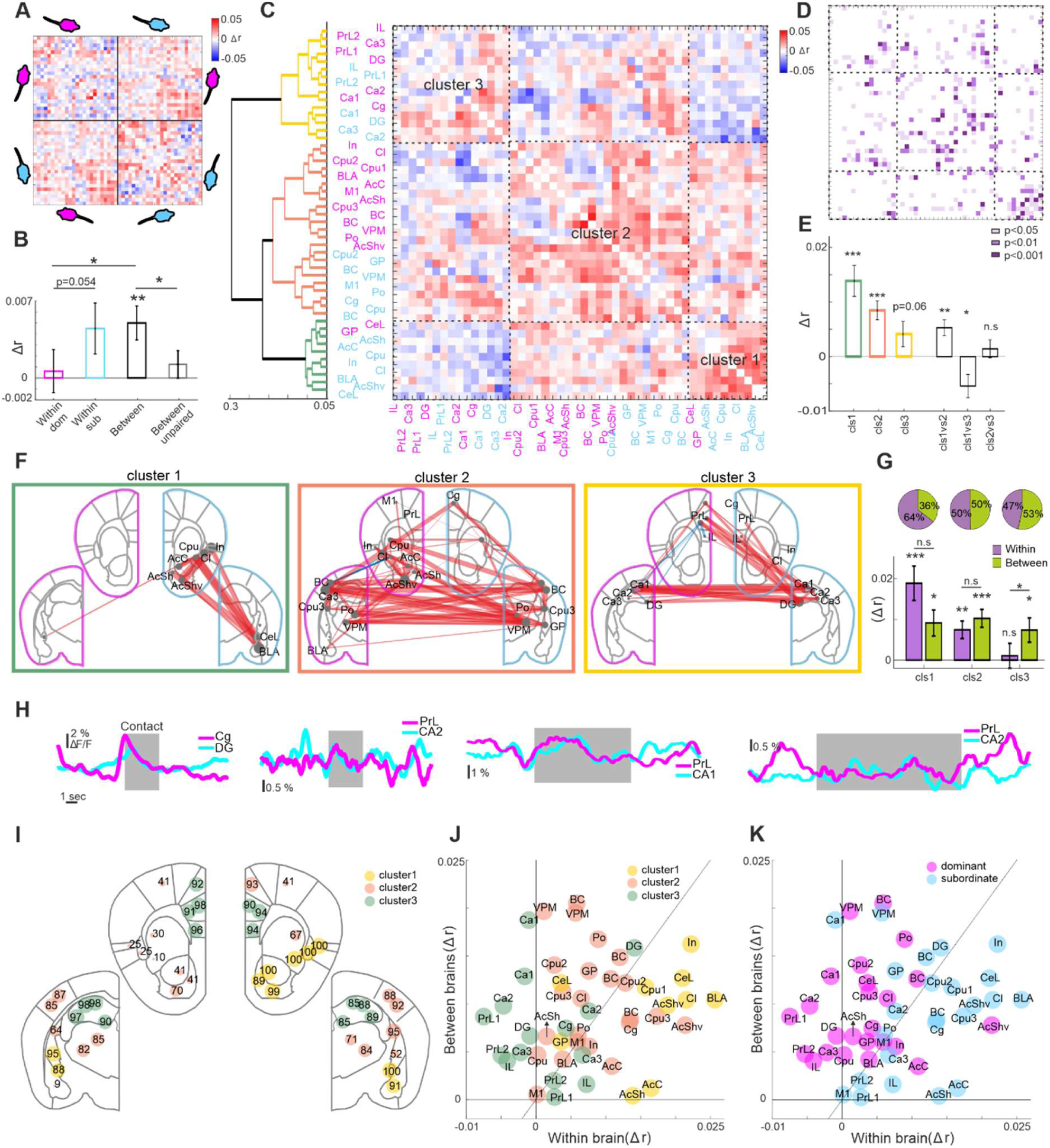
Brain-wide subnetworks during social interactions. **a.** Differential correlation matrix (correlation during contact minus correlation during no-contact; Δr) between all brain areas (pair-wise) within and between two mice. Each element in the matrix depicts the Δr value between a pair of brain areas. Red colors indicate a bias for contact and blue colors indicate a bias for no contact periods. Matrix averages over 42 pairs of mice in which the dominant mouse is always on the top left quadrant (pink) and the subordinate mouse is always on the bottom right quadrant (cyan). **b.** Grand average Δr values within brain areas of the dominant (pink), subordinate (cyan) and between mice (black). An unpaired control (correlation between shuffled pairs) is presented in gray. Error bars depict mean±SEM across pairs (n=42). **c.** Differential correlation matrix from a, now sorted based on hierarchical clustering (Ward’s method; Dendrogram on the left) identifying three distinct clusters (colored on the dendrogram). Brain areas are colored based on dominant (pink) or subordinate (cyan) mouse. **d.** Significance matrix corresponding to the matrix in c in which significant pairs ( Wilcoxon signed-rank test) are color coded for significance level. **e.** Δr values in each cluster and between clusters. Error bars as in b. **f.** Δr values superimposed on coronal sections for each cluster separately. Width of each line depict Δr magnitude. Red lines indicate a bias towards contact whereas blue lines indicate a bias towards no-contact. Only 30% extreme Δr values are shown. **g.** Average Δr values for each cluster divided to within (purple) and between (green) connections. Error bars as in b. Proportion of within and between connections are on top. **h.** Example traces of a prefrontal response in the dominant mouse (pink) and a simultaneous hippocampal response in the subordinate mouse (cyan). Shaded gray areas depict social interaction. **i.** Percentage of reproducibility for the assignment of each brain area to its cluster (Bootstrap analysis; see Methods). High percentage indicates stability of that brain area to its cluster (in f). **j.** Δr values between brains versus within brain for each brain area separately. Different clusters are color coded. **k.** Same as in j but color coded for dominant and subordinate mice. * - *p*<0.05, ** - *p*<0.01, *** - *p*<0.001, Wilcoxon signed-rank test.

The differential correlation matrix in Fig. 2a revealed structured patterns suggestive of functional subnetworks across and within brains. Using hierarchical clustering we sorted the differential correlation matrix based on similarity (Ward’s method) and obtained three distinct clusters, i.e., subnetworks (Fig. 2c). Interestingly, each cluster contains brain areas belonging to both the dominant (pink) and subordinate (cyan) mice and both within and between brains connections. Pairwise statistical testing showed a high density of significant Δr values within clusters (Fig. 2d). Δr values within clusters 1 and 2 were significantly greater than zero (*p<*0.001; Wilcoxon signed-rank test, n=42) with a positive trend also in cluster 3 (p=0.06; Fig. 2e). In addition, Δr values between clusters have a significant positive difference between clusters 1 and 2 (*p*<0.01; Wilcoxon signed-rank test), a significant negative difference between clusters 1 and 3 (*p*<0.05) and a non-significant difference between clusters 2 and 3 (*p*>0.05). A comparable clustering structure was recovered using an alternative method based on t-SNE projections (Fig. S4. b), supporting the robustness of the results.

To visualize each subnetwork (i.e., cluster) we superimposed the top 30% of Δr values within each cluster onto brain outlines (Fig. 2f). Red lines indicate a bias towards social contact, while blue lines reflect a bias towards no-contact epochs. The dominant mouse (pink) is shown on the left, and the subordinate (cyan) on the right. This representation revealed that cluster 1 consists primarily of amygdala and nucleus accumbens in the subordinate brain. Cluster 2 includes sensory-related areas from both mice spanning thalamus and cortex. Cluster 3 encompasses prefrontal cortex and the hippocampus from both mice with a notable bias in connections from the dominant prefrontal cortex to the hippocampus of the subordinate. To further characterize these subnetworks, we quantified within and between brains Δr values within each cluster (Fig. 2g). Cluster 1 had more within brain connections (64%) and significantly positive Δr values within brains (*p*<0.001; Wilcoxon signed-rank test n=42) with smaller but still significant between brains Δr values (*p*<0.05). Cluster 2 had an equal proportion of within- and between-brain connections, and both were significantly positive (*p*<0.01). Cluster 3 showed slightly more connections between brains (53%) and significantly positive Δr values only between brains, with a significant difference compared to within-brain Δr values (*p*<0.05). To evaluate interactions between subnetworks, we examined Δr values between clusters (1 vs. 2, 1 vs. 3, 2 vs. 3), revealing several significant differences (Fig. S4. d,e). Representative traces from the dominant prefrontal cortex and subordinate hippocampus illustrate this increased coupling during social contact (Fig. 2h). To assess the stability of the observed clusters, we performed a bootstrap analysis by randomly selecting 80% of mice pairs and repeating the clustering procedure over 1,000 iterations. For each iteration, the cluster identity of each brain area was matched to its original cluster. Most regions displayed stability in terms of cluster identity (>80%), particularly those with strong and central connectivity (Fig. 2i). Some regions such as the BLA of the dominant mouse showed low stability (9%), suggesting they may belong to an alternative subnetwork (see Methods). The full distribution of cluster assignments across all brain areas is provided in Figure. S4. f.

As in Fig. 1, we performed several controls to ensure the observed subnetwork structure was specific to social contact: (1) We randomly split ‘no contact’ epochs into two groups, computed the differential correlation matrix, followed by clustering. This yielded a shallower dendrogram (Fig. S5. b) with mostly non-significant Δr values within each cluster (*p*>0.05; Wilcoxon signed-rank test; Fig. S5. c). (2) To control for possible common input effects, we applied the unpaired control used in Fig. 2b, shuffling contact epochs between mice, and again observed a shallow dendrogram with no significant within-cluster Δr values (*p* > 0.05; Fig. S5. e-h).

Next, we quantified the relationship between and within brains or each brain area separately. To do this, we plotted the Δr values for each brain area (within its own cluster) averaged within the brain against the average between brains. When color-coded by cluster, cluster 1 exhibited a bias toward within-brain connectivity, cluster 3 toward between-brain connectivity, and cluster 2 showed no consistent bias (Fig. 2j). When color-coded by dominance, brain areas in dominant mice tended to favor between-brain connections, whereas subordinate brain areas were biased toward within-brain connectivity (Fig. 2k). In summary, the network analysis revealed distinct brain-wide subnetworks that emerge during social interaction and uniquely span within and between brains.

### Social rank asymmetries within and between brains

As shown in Fig. 2, several connections exhibit a clear asymmetry between dominant and subordinate mice. To quantify asymmetry, we defined an asymmetry index (ASI) for each edge (i.e., Δr between a pair of brain regions) by subtracting the Δr of the dominant mouse from the Δr of the subordinate mouse divided by their absolute sum (Fig. 3a). This can be done for within and between brain connections. The ASI ranges from −1 to 1, where positive values indicate a bias towards the dominant mouse and negative values indicate a bias towards the subordinate mouse. Overall, ASI values were generally negative within brains and positive between brains, indicating that during social contact, the dominant brain shows greater coupling with the subordinate brain, while the subordinate brain shows increased internal synchrony (Fig. 3b). Among subnetworks, cluster 3 (i.e., prefrontal hippocampus subnetwork), exhibited the strongest asymmetry, with significantly positive ASI between brains (*p* < 0.01) and significantly negative ASI within brains (*p* < 0.05, Wilcoxon signed-rank test, *n* = 42; Fig. 3c). Cluster 2 also showed significantly positive ASI between brains (*p*<0.001). Sorting ASI values across connections revealed specific motifs: prefrontal-hippocampus connections in cluster 3 showed high ASI between brains but negative values within brains (Fig. 3d), while the cingulate–barrel cortex connection in cluster 2 showed high inter-brain ASI. These results suggest that, during social interaction, the dominant mouse’s prefrontal cortex preferentially couples with the subordinate’s hippocampus whereas internal pathways in the subordinate mouse increase.

**Figure 3.**
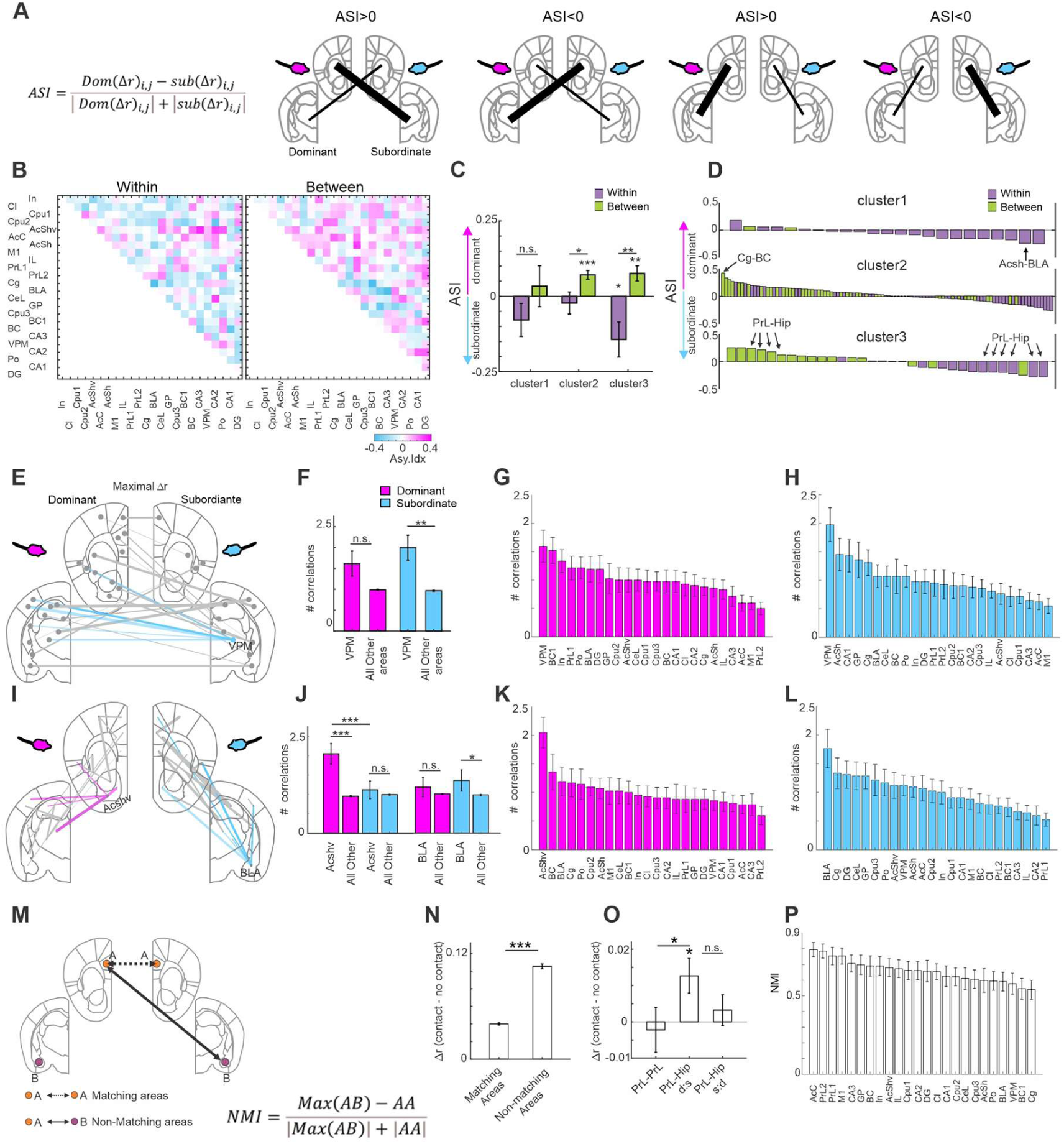
Social rank asymmetries within brain-wide subnetworks. **a.** Schematic illustration and equation for the asymmetry index (ASI). **b.** Asymmetry index (ASI) for social rank in each connection within (left) or between (right) brains. Pink color indicates a bias for the dominant mouse whereas cyan colors indicate a bias for the subordinate mouse. Average across 42 pairs. **c.** ASI for each cluster divided for within (purple) and between (green) brains. Error bars depict mean±SEM across pairs (n=42). **d.** ASI sorted pairs for each cluster for within (purple) and between (green) connections. **e.** Plots of the maximal Δr between brains in the dominant mouse. Each brain area in the dominant mouse has one connection. Connections with the VPM of the subordinate mouse are colored in cyan. **f.** The number of maximal connections in the VPM vs all other brain areas in the subordinate (cyan) and dominant (pink) mouse. Error bars as in c. **g, h.** Number of maximal connections sorted across brain areas in the dominant and subordinate mouse. **i-l.** Same as f-h but for within brains. **m.** A schematic illustrating matching (homologous) versus non-matching inter-brain connections. **n.** Δr values between matching areas and the maximal Δr between non-matching areas. **o.** Specific example for Δr values between both prelimbic cortices (matching) and prelimbic-hippocampus alternating between dominant and subordinate mice. Error bars as in c. **p.** Non-matching index sorted across brain areas (see Methods). * - *p*<0.05, ** - *p*<0.01, *** - *p*<0.001, Wilcoxon signed-rank test.

To identify central brain regions that may serve as hubs, i.e., nodes with strong connections to many other regions, we isolated, for each brain area, the single strongest Δr connection (in absolute value) with all other areas. This was computed separately for within- and between-brain connections. Between brains, the VPM in the subordinate mouse showed the highest number of connections with the dominant mouse (Fig. 3e). The number of connections with the VPM of the subordinate mouse was significantly greater than the number of connections of other areas (*p*<0.01; Wilcoxon signed-rank test), but not significant in the dominant mouse (*p*>0.05; Fig. 3f). Sorting regions by the number of maximal connections further emphasized this (Fig. 3g,h). Within brains, the nucleus accumbens emerged as the most central hub in the dominant mouse, while the basolateral amygdala (BLA) was most central in the subordinate mouse (Fig. 3i-l). Together, during social contact, the VPM and the BLA of the subordinate mouse and the accumbens of the dominant mouse displayed hub dynamics.

Our dual-mouse brain-wide approach, enables measurement of inter-brain dynamics not limited to anatomically homologous regions in contrast to prior studies that focused primarily on same-area interactions, for example prefrontal-prefrontal (Fig. 3m;(*5*, *8*, *9*, *27*)). We found that maximal Δr values between non-matching areas were significantly higher than the Δr values between matching (homologous) areas (*p*<0.001; Wilcoxon signed-rank test; Fig. 3n). In particular, correlations between the prelimbic cortex of the dominant mouse and the hippocampus of the subordinate were significantly stronger than between the two prelimbic cortices (*p*<0.05; Fig. 3o). To quantify this effect across the brain, we computed a non-matching index (NMI; i.e., Δr for strongest non-matching region minus Δr for the homologous region) and sorted across regions (Fig. 3p). Prelimbic areas showed high NMI values but also many other areas displayed positive values (Fig. 3p). In summary, we find that brain areas of one mouse interact stronger with different brain areas of the other mouse, not necessarily the homologous area.

### Prefrontal cortex of the dominant mouse is enhanced before social interaction

To investigate whether meaningful brain activity occurs before social interaction, we grouped mouse pairs based on social hierarchy into high rank ratio pairs (*n* = 20) and equal rank ratio pairs (*n* = 22; Fig. 4a; see Methods). We focused on interactions initiated by the higher-ranked mouse. Neural responses were aligned to social contact and plotted for each brain region, grouped by cluster and rank condition (Fig. 4b). Overall, high rank pairs exhibited distinct neural dynamics compared to equal rank pairs. We focused on three key brain structures: (1) The prefrontal cortex (PrL, IL, Cg pooled together) showed significantly elevated response in the higher ranked mouse before the social contact and only in high rank pairs, possibly reflecting dominance excretion over subordinate prior to engagement (Fig. 4c, d; p<0.01; Wilcoxon signed-rank test). (2) The amygdala (BLA and CeL pooled together) showed significantly elevated response in the lower ranked mouse around the social contact and only in high rank pairs, possibly reflecting a fear or threat-related response (Fig. 4c, d; *p*<0.01; Wilcoxon signed-rank test). (3) The hippocampus (grouped across hippocampal areas) showed a significantly higher response in the lower ranked mouse after social contact and only in high rank pairs, potentially linked to updating social memory or reinforcing hierarchical structure (Fig. 4c, d; *p*<0.01; Wilcoxon signed-rank test).

**Figure 4.**
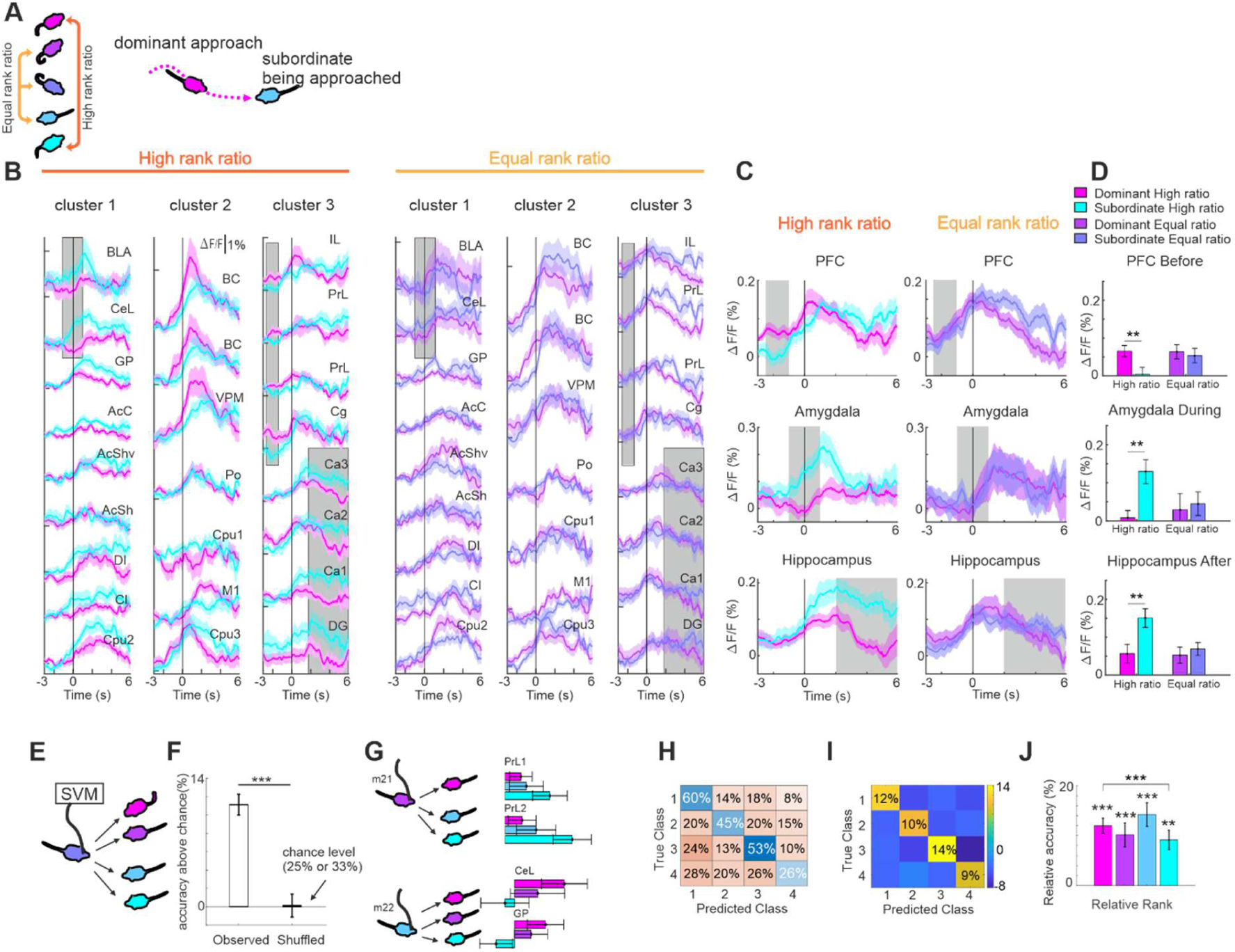
Prefrontal cortex of the dominant mouse encodes initiation of a social interaction. **a.** Schematic of high and equal rank ratio pairs and focusing on interaction upon which the higher ranked mouse initiated social interaction. **b**. Neuronal responses in 24 brain areas (divided into clusters) for high and equal rank pairs. **c.** Averaged traces of high (left) and equal (right) rank in prefrontal, amygdala and hippocampus. Time periods of interest (before, during and after social contact) are marked in gray shades. **d.** Average responses for high and equal rank pairs averaged within specific time epochs (gray shades in c) for each area presented in c. Error bars depict mean±SEM across mice (20 high rank and 22 equal rank). **e.** Schematic illustration for an SVM classifier that predicts the identity of the partner mouse based on the brain of one mouse. **f.** Accuracy (% above chance) of the identity classifier for observed and trial shuffled data. Error bars depict mean± SEM across mice (n=23). **g.** Example responses from one mouse when interacting with different mice. Error bars depict mean±SEM across social interactions (m21: n=356, 281, 308 from dominant to subordinate, respectively; m22: n = 364, 344, 281) **h.** Example confusion matrix from one mouse, indicating a detailed accuracy profile for each mouse identity (1 = dominant, 4 = subordinate). **i.** Average normalized confusion matrix (observed accuracy minus shuffled accuracy for each cell) across mice. **j.** Average accuracy for each mouse rank. Error bars as in f. * - *p*<0.05, ** - *p*<0.01, *** - *p*<0.001, Wilcoxon signed-rank test.

Continuing this line, we trained a classifier to predict whether a mouse is an initiator, based on brain responses (SVM; see Methods). This was done on epochs before, during or after social contact (Fig. S6. a). During all time epochs the classifier was significantly higher than a trial shuffle in both high and equal rank with a significantly higher accuracy in high rank compared to equal rank pairs (*p*<0.001; 74% vs 62%; Wilcoxon Rank-sum test; Fig. S6. a). Observing the weights of the classifier, positive weights indicate a higher activity in the initiator, whereas negative weights indicate higher activity in the mouse being approached. Before contact prefrontal areas had positive weights, during contact sensory areas had positive weights and amygdala showed negative weights and after contact, hippocampal regions displayed negative weights (Fig. S6. b). Together, these results suggest a temporal cascade of activity during social interaction: prefrontal activation in the initiator precedes amygdala activation in the subordinate along with mutual sensation in both mice, followed by hippocampal responses that may reflect social memory or evaluation.

Finally, we tested whether a mouse’s brain-wide activity during social contact encodes information about the identity of its interaction partner. To this end, we trained a support vector machine (SVM) classifier to predict partner identity (i.e., 3 or 4 partners within its group) using the subject’s own neural activity during contact epochs (Fig. 4e; see Methods). Classifier accuracy was significantly higher compared to trial shuffled data (12.0% ± 1.24% above chance, mean ± SEM; *p*<0.001; Wilcoxon signed-rank test; Fig. 4f). Example responses from one mouse when interacting with different partners shows a ranked response profile in areas such as prelimbic and amygdala (Fig. 4g). An example confusion matrix (Fig. 4h) and the average confusion matrix (Fig. 4i) indicates the ability of the classifier to accurately predict across all partners. Each identity class was predicted significantly above chance (*p* < 0.001), with accuracy significantly higher for dominant partners compared to subordinates (*p* < 0.001; Fig. 4j). Together, these results indicate that brain-wide dynamics also encode partner identity-an essential feature of naturalistic social behavior.

### ASD-like mice exhibit minimal inter-brain synchrony and enhanced within brain synchrony

The results above provide a comprehensive quantification of brain-wide subnetworks during social interaction in normally behaving wild-type mice. We next asked whether these dynamics are altered in animals with social impairments. To this end, we performed additional experiments in a Shank3 mouse model of autism spectrum disorder (ASD mice). Multi-fiber arrays were implanted into 24 mice from 7 social groups, including ASD-only groups (n = 2) and mixed groups containing both ASD and WT animals (n = 5; Fig. S7. a). ASD mice were tested under identical naturalistic conditions as WT controls. Behaviorally, ASD mice engaged in significantly fewer social interactions (Fig. S7. b; *p*<0.001; Wilcoxon Rank-sum test; n=48 and 34 pairs in WT and ASD mice respectively) and significantly less prosocial behaviors such as ‘nose-to-nose’ and ‘side-by-side’ interactions (Fig. S7c; *p*<0.001; Wilcoxon Rank-sum test). Instead, they spent significantly more time ‘sitting together’ (Fig. S7. c) and exhibited deficits in additional behavioral assays (Fig. S7. e). Within mixed groups, ASD mice consistently occupied lower social ranks, whereas their rank distribution in ASD-only pairs was broader and comparable to that of WT animals (Fig. S7. d). These relative social ranks were correlated to other behavioral parameters (Fig. S7. f-j). At the neural level, aligning activity to social contact revealed a widespread enhancement in ASD mice, with significant increases across all recorded brain regions (Fig. 5a-c; *p*<0.01; Signed rank test with Bonferroni correction; n=24). A classifier trained on these signals decoded social contact with high accuracy (88%±6% compared to shuffle 50%±2%) at levels comparable to WT mice. Compared to WT, responses were significantly elevated in several regions: barrel cortex (BC), anterior cingulate cortex (ACC), cingulate cortex (Cg), and dentate gyrus (DG) (Fig. 5d; *p*<0.05; Wilcoxon Rank-Sum test with FDR correction; See Fig. S8 for all brain areas). In summary, ASD mice displayed impaired social interactions accompanied with enhanced brain-wide responses around social contact.

**Figure 5.**
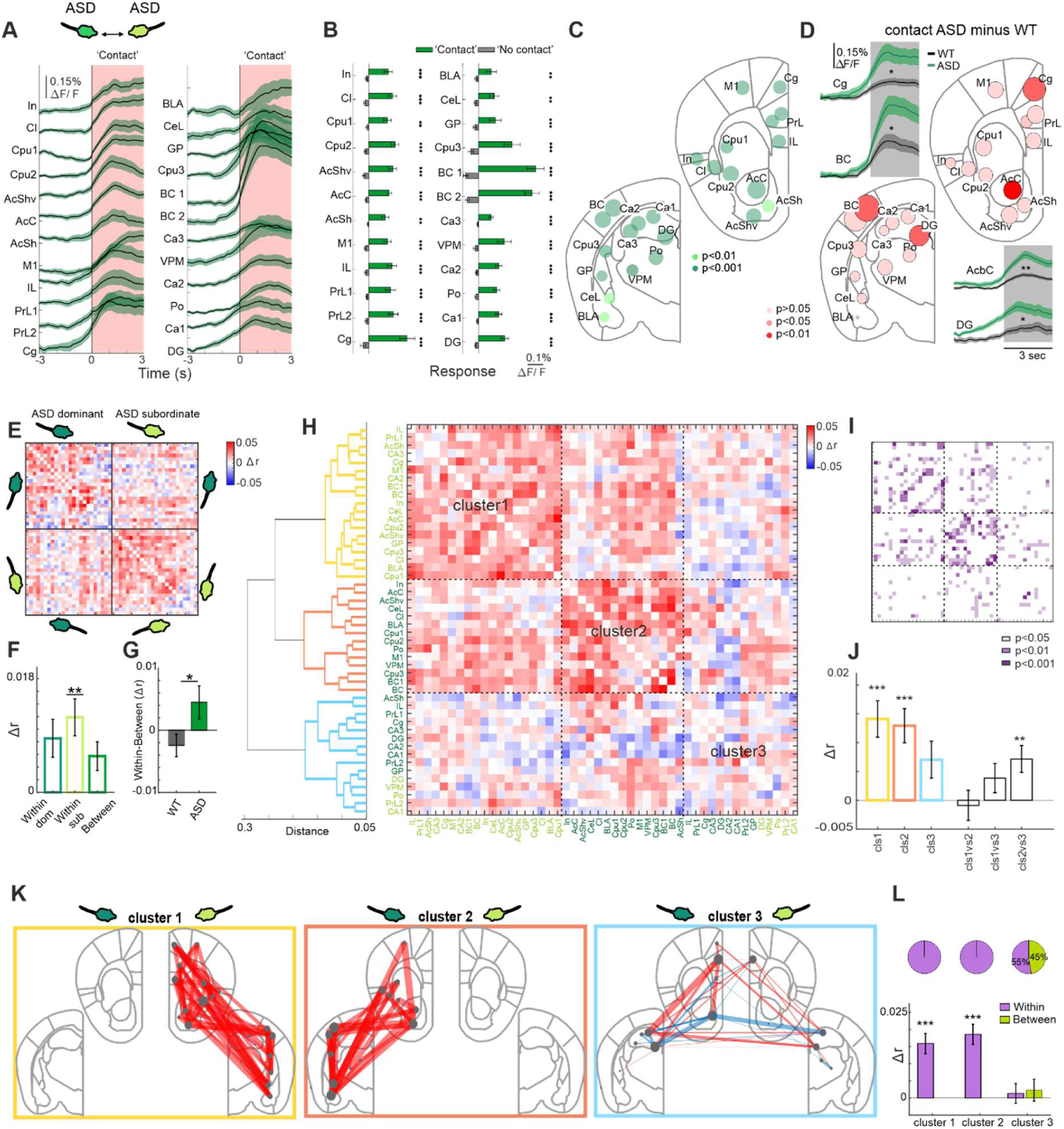
ASD mice display brain-wide hyper-activity along with enhanced within brain synchrony. **a.** Responses (ΔF/F) in 24 brain areas aligned on social contact (time 0) for ASD mice. Error bars depict mean±SEM across mice (n=24). **b.** Grand average responses of ASD mice averaged during social interaction (i.e., contact; green) and outside social interaction (i.e., no-contact; gray). Error bras as in a. *** *p* <0.001,\*\**p*<0.01,\**p*<0.05, Wilcoxon signed test with Bonferroni correction (n=24). **c.** Circle representation depicting the difference in responses between contact and no-contact periods divided by the sum the SEM of both. The larger the circle the bigger the difference between contact and no-contact activation. Significance levels are color coded. **d.** Circle representations depicting the difference in responses between ASD and WT during social contact (0-3 seconds after contact). The circle size denote the differences between ASD and WT mice and the color, the significance level. Significant responses in specific areas for ASD (green) and WT (black) are additionally plotted. * - *p*<0.05; ** - *p*<0.01; Wilcoxon sum rank test; FDR corrected. **e.** Differential correlation matrix (correlation during contact minus correlation during no-contact; Δr) between all brain areas (pair-wise) within and between two mice. Each element in the matrix depicts the Δr value between a pair of brain areas. Red colors indicate a bias for contact and blue colors indicate a bias for no contact periods. Matrix averages over 34 pairs of mice in which the dominant mouse is always on the top left quadrant (dark green) and the subordinate mouse is always on the bottom right quadrant (light green). **f.** Grand average Δr values within brain areas of the dominant (dark green), subordinate (light green) and between mice (green). Error bars depict mean±SEM across pairs (n=34). **g.** Difference of within Δr minus between Δr in WT (gray) and ASD (green) mice. *-p<0.05 Wilcoxon rank-sum test. **h.** Differential correlation matrix from a, now sorted based on hierarchical clustering (Ward’s method; Dendrogram on the left) identifying three distinct clusters (colored on the dendrogram). Brain areas are colored based on dominant (dark green) or subordinate (light green) mouse. **i.** Significance matrix corresponding to the matrix in d in which significant pairs (Wilcoxon signed-rank test) are color coded for significance level. **j.** Δr values in each cluster and between clusters. Error bars as in b. **k.** Δr values superimposed on coronal sections for each cluster separately. Width of each line depict Δr magnitude. Red lines indicate a bias towards contact whereas blue lines indicate a bias towards no-contact. Only 30% extreme Δr values are shown. **l.** Average Δr values for each cluster divided to within (purple) and between (green) connections. Error bars as in f. Proportion of within and between connections are on top. * - *p*<0.05, ** - *p*<0.01, *** - *p*<0.001, Wilcoxon signed-rank test.

Next, we investigated brain-wide subnetworks within and between ASD mice. Using the same analysis applied to WT animals (Fig. 2) we computed the differential correlation matrix (Δr; contact minus non-contact) for all ASD pairs, preserving dominance order (Fig. 5e; top left is the dominant ASD mouse; Bottom right is the subordinate ASD mouse). ASD mice exhibited elevated within-brain Δr values, significant for the subordinate ASD mouse (*p*<0.01; Wilcoxon Signed rank test) but reduced Δr values between brains (Fig. 5f). The within–between difference was significantly larger in ASD than WT mice, indicating an opposite ratio, biased toward within-brain synchrony (Fig. 5g; *p*<0.05; Wilcoxon sum rank test). Clustering of the differential correlation matrix revealed three subnetworks (Fig. 5h), Two clusters contained a substantial number of significant connections (non-zero Δr) and showed robustly positive average Δr values, whereas the third cluster was not significantly different from zero (Fig. 5i, j). Interestingly, when mapped onto anatomical outlines (top 30% Δr values within each cluster), the two significantly positive clusters map exclusively to each ASD mouse separately with no inter-brain connection (Fig. 5k,l). In contrast, the third nonsignificant cluster showed weak inter-brain synchrony between prefrontal and hippocampus, resembling the strong prefrontal/hippocampus cluster in WT mice. To quantify this, we divided each cluster into within and between connections (Δr values) and found significantly within brain Δr values for the two clusters (*p*<0.001; Wilcoxon signed-rank test) and nonexistent between Δr values. Cluster 3 did not display significant Δr values for both within and between connections. In summary, ASD mice during social contact correlated response within their own brain with a minor inter-brain synchronization.

Finally, we studied the temporal dynamics of brain-wide responses around social contact in ASD mice. As in WT animals, we divided pairs according to rank ratio and focused on interactions in which the dominant mouse initiated contact (Fig. 6a; n=19, and 15 pairs high rank and equal rank, respectively; note that the ASD pairs displayed an even greater rank ratio than WT pairs). Unlike WT mice, dominant ASD animals did not exhibit significant elevated prefrontal activity before initiating contact (Fig. 6b,c; for both high and equal rank pairs; *p*>0.05; Wilcoxon Signed Rank test). In addition, amygdala responses were not significantly modulated during contact (*p* > 0.05), although subordinate ASD mice in high-rank pairs showed a delayed enhancement shortly after contact. Hippocampal responses also failed to show the post-contact increase seen in WT subordinates (*p* > 0.05 for both rank conditions). Overall, the temporal dynamics within ASD pairs were less discriminative around social contact, as compared to WT mice. The full temporal dynamics in each area is plotted in Fig. S9. Consistent with this, an SVM classifier trained on brain-wide dynamics predicted the partner’s identity with lower accuracy in ASD than in WT mice (9.5% vs. 11.5% above shuffle; *p* = 0.07, Wilcoxon rank-sum test), hinting of an impaired theory of mind in ASD mice (see discussion). Together, these results suggest that while ASD mice exhibit strong within-brain subnetworks, they display minor dynamics related to the partner mouse, potentially reflecting an impaired capacity for modeling the partner’s brain state.

**Figure 6.**
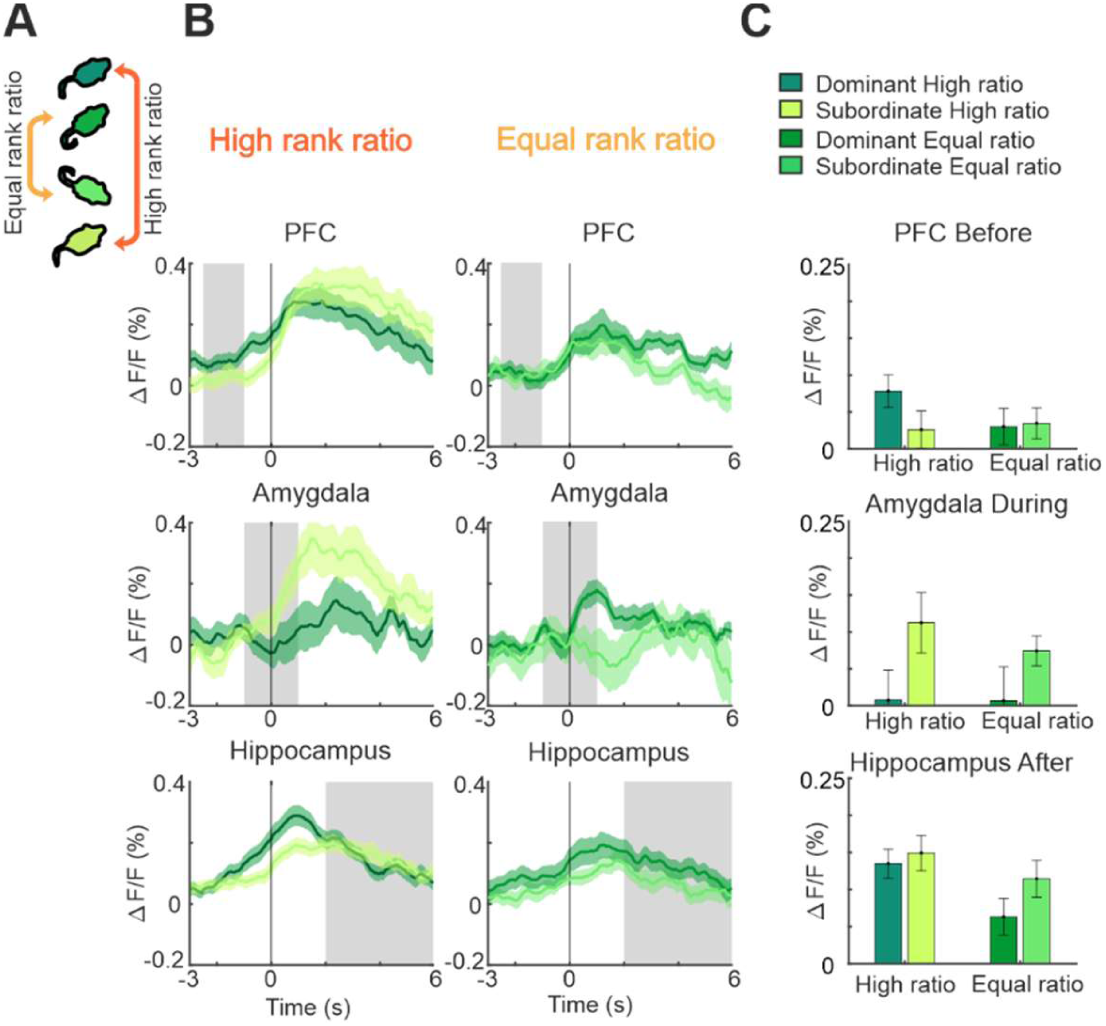
ASD mice display minor differences in temporal dynamics. **a.** Schematic of high and equal rank ratio pairs **b.** averaged traces of high (left) and equal (right) rank in prefrontal, amygdala and hippocampus. Time periods of interest (before, during and after social contact) are marked in gray shades. **c.** Average responses for high and equal rank pairs averaged within specific time epochs (gray shades in b) for each area presented in b. Error bars depict mean±SEM across mice (19 high rank and 15 equal rank).

## Discussion

### The social brain is widespread

This study explores brain-wide dynamics during social interactions in a more naturalistic setting as compared to previous studies using confined social tests such as a three-chamber test or a tube test (*6*, *10*, *58*, *19–21*, *53–57*). Naturalistic interactions elicited widespread activation not only in canonical social regions (such as amygdala, prefrontal cortex and nucleus accumbens;(*35*, *59–61*)) but also in areas not traditionally social-related (such as VPM, Po in thalamus, BC and M1 in cortex and to some extent striatum and hippocampus; (*62*, *63*)). From a holistic viewpoint, social interaction between two mice engages mutual whisking, multimodal sensory integration, movement and encoding of partner identity, interaction type, and affective context, processes likely engaging broad neural circuits. Notably, such brain-wide responses were absent during control conditions involving object contact or movement alone, underscoring the social specificity of the observed activity. In addition, interactions within familiar cage mates may cause a higher and broader response profile as compared to stranger mice(*9*, *13*, *64*) and natural interactions elicit stronger responses than interactions separated by a mesh (*65*). Despite this global engagement, distinct brain areas showed distinct temporal profiles, for instance, some areas such as the prefrontal cortex and hippocampus have an early (before social contact) response onset whereas other areas respond later. These differences suggest complex, temporally structured dynamics within the brain-wide network, that merit further dissection.

### Encoding of social information is interwoven between the two brains

In this study we were able to simultaneously measure brain-wide dynamics from two freely interacting mice, enabling the investigation of inter-brain dynamics on a broad scale. By measuring from all pairs within a group we inferred relative social ranks and examined how brain-wide interactions vary with dominance hierarchy. Our network analysis revealed three distinct subnetworks: (1) an amygdala-accumbens subnetwork present specifically in subordinate mice(*66*, *67*). This subnetwork also included the claustrum and the insular cortex which are also relevant in an emotional context as well as during multimodal integration(*68–74*). We call this the emotional subnetwork and emphasize its relation to negative or passive social contexts in which one is being approached and possibly facing forceful behavior(*57*, *65*, *75–80*). The emotional subnetwork may display a chronic over-activity that is present in subjects with high anxiety, e.g., social anxiety. (2) A thalamus-cortex-striatum subnetwork equally distributed across dominant and subordinate mice and involves both within and between brain interactions. We call this the sensory subnetwork which may encode sensory integration during social contact (*62*, *63*, *81–84*). We highlight that dominant mice display higher inter-brain correlations within this subnetwork (Fig. 3c) maybe because of their initiating role and need to precisely perceive the situation. Conversely, the VPM thalamus of the subordinate mouse receives the most interactions from the brain of the dominant mouse (Fig. 3e,f), indicating its role in sensory alertness when being approached by another. (3) A prefrontal-hippocampus subnetwork which is biased toward inter-brain correlations (Fig. 2f,g) where each region is known to encoding social information(*8*, *13*, *90–94*, *21*, *54*, *56*, *85–89*) and the hippocampal-mPFC pathway within the brain encodes social memory(*94*, *95*). We call this the decision/consolidation subnetwork and highlight a specific asymmetry between the prefrontal and hippocampus which is biased towards the dominant mouse (Fig. 2f, Fig. 3,b,c,o). We propose that dominants initiate interactions via prefrontal activation, whereas subordinates consolidate the experience via hippocampus (Fig. 4c,d).

Furthermore, we found a temporal profile along the social interaction, starting with activity in the prefrontal of the dominant well before social contact, followed by a mutual activation of the emotional and sensory subnetworks around social onset and ending with hippocampal activity in the subordinate mouse (Fig. 4b-d). This timeline further supports the logic of the interplay between the different subnetworks. Several connections within each subnetwork are indeed related to anatomical connectivity (e.g., between amygdala and accumbens(*66*, *67*); or prefrontal and hippocampus(*96–100*)), but we highlight that many connections are across brains making them irrelevant to within brain anatomy. In summary, we identify three interwoven subnetworks, emotional, sensory, and decision/consolidation, distributed across brains and coordinated in time, which together may underlie the neural encoding of natural social interaction.

### Prefrontal-hippocampus interactions across brains

Brain-wide subnetworks differed by social rank. Dominant mice exhibited stronger inter-brain synchrony, even greater than their intra-brain synchrony (Fig. 2b), suggesting an active role in sensing and engaging the partner. This was especially salient within the decision/consolidation subnetwork in which the prefrontal cortex of the dominant mouse was correlated with hippocampus of the subordinate (Fig. 2f, Fig. 3,b,c,o). The prefrontal cortex was found to be particularly important in the dominant mouse displaying enhanced response and predictive power seconds before social contact. Notably, this anticipatory prefrontal activity was attenuated in interactions between similarly ranked mice, where social contact appeared more mutual and less hierarchical (Fig. 4b-d). These findings align with previous studies showing that prefrontal cortex encodes higher-order social information and is related to social rank(*6*, *21*, *54*, *56*, *85–88*, *101*), including prior response(*101*) and specific synchrony between high rank ratio pairs(*8*).

In contrast, the subordinate mouse is mostly being approached and is more anxious, activating its own emotional network including the amygdala (Fig. 2f), particularly when approached by a substantially higher-ranked partner (Fig. 4c,d). Amygdala enhancement and predictive power around the social touch is in line with previous findings relating amygdala to social information with emphasis on negative experience(*78*, *79*, *102–104*). Following social contact with a highly dominant mouse, the subordinate hippocampus showed a marked increase in activity. This may reflect the encoding of social memory and consolidation of specific social information such as identity and rank(*13*, *87*, *90*, *91*, *105*). Although the dominant hippocampus also responded during interaction, the effect was weaker (Fig. 4c), suggesting that social memory consolidation occurs in both partners but is more robust when the interaction is imposed rather than initiated. Together, we highlight an asymmetric interaction between dominant prefrontal cortex and the subordinate hippocampus, underscoring the importance of studying inter-brain dynamics across non-homologous regions which was previously largely unexplored. Our multi-area dual brain approach further enabled us to observe hub regions in one mouse that were connected to several brain regions in the partner mouse, for example the VPM of the subordinate mouse, which is considered a sensory relay station to the cortex. Thus, our innovative approach adds novel and crucial insights to the field of social neuroscience, conceptually needing two brains to fully understand social interaction.

### The ASD-like brain globally interacts within itself, not with the partner brain

To investigate how brain-wide networks operate under socially abnormal scenarios, we recorded large-scale neural activity in ASD mouse model during natural social interactions. Contrary to much of the existing literature in both humans (*106–111*)and mice (*48–50*, *112–115*), which reports reduced activation in key regions such as the amygdala, prefrontal cortex, and hippocampus, we observed a striking brain-wide hyperactivation in ASD mice, often exceeding that of wild-type controls (consistent with (*45*, *116*)). This discrepancy may stem from several factors, including differences in mouse lines, recording methods, brain regions examined, or behavioral paradigms. Importantly, our paradigm captures naturalistic, freely moving social interactions, which may be more cognitively and emotionally demanding for ASD mice, potentially eliciting a global increase in neural activity. Certain regions, such as the barrel cortex (BC) and dentate gyrus (DG), may act in a compensatory capacity, attempting to stabilize dysfunctional processing elsewhere in the network (*108*, *117*, *118*). Alternatively, the overactivation in ASD mice can be linked to oversensitivity to external stimuli present in ASD individuals and may lead to social avoidance (*45*, *119–122*). Additionally, hyperactivity may reflect an altered excitation/inhibition (E/I) balance (but not necessarily (*123*)), where impaired inhibition, possibly due to reduced parvalbumin-positive (PV⁺) interneuron function, shifts the local circuitry toward excess excitation (*123–127*). While some studies support elevated E/I ratios in areas like the prefrontal and somatosensory cortices(*123–126*, *128*, *129*), others report heterogeneous patterns across brain areas (*130*). Whether a unifying circuit mechanism underlies these global changes remains an open question for future investigation.

Our data further reveals that during social interaction, ASD mice display strong within-brain synchrony but minimal inter-brain synchrony. While a decision/consolidation subnetwork is still detectable, its expression is notably weak. These results suggest that the autistic brain may exhibit a kind of functional insularity, engaging in overactive, self-referential processing that inhibits alignment with a social partner. This internal hyper-coherence could underlie behavioral symptoms such as difficulty with eye contact or reciprocal social cues, as the neural network may be too inwardly focused to coordinate across individuals.

### Pitfalls and future directions

The current study is focused on network-level dynamics within and between brains, measuring bulk non-specific neuronal populations. But social information could further be encoded within specific neuronal subtypes, for example inhibitory neurons(*5*, *23*, *131*), projection specific neurons(*10*, *21*, *56*, *132*) or even non-neuronal cells(*55*). Network-level dynamics could also be affected by neuromodulatory signals, such as dopamine, serotonine, oxytocin and others(*58*, *92*, *133*, *134*), which may underlie social rank biases within and between brains and whose dysregulation has been linked to social deficits in ASD (*135–142*). Notably, our optical approach enables us to investigate network-level dynamics of specific neuronal subtypes and neuromodulatory signals that can complement our findings. Moreover, the multi-fiber photometry platform can be readily adapted to target additional brain areas not covered here, such as the hypothalamus(*21*, *143*) orbitofrontal cortex(*144*, *145*) and other brain areas that are not classically associated with social engagement. As a complementary method, freely moving wide-field imaging(*146*) could capture cortex-wide dynamics during naturalistic social interaction, extending our knowledge beyond well studied head-fixed behaviors(*147–149*). Future studies could probe causal effects at the network level through patterned optogenetics to silence or activate specific subnetworks within certain individuals, and may rescue abnormal social behaviors in ASD mice.

An interesting finding is that brain-wide dynamics can predict the identity of the partner mouse, a key element of social engagement. This finding provides direct support for the theory of mind (ToM) hypothesis that describes the cognitive ability to infer the mental states of others, a crucial aspect of social interaction enabling to interpret the partner’s behavior(*150*). Impairments in ToM have been proposed to underlie social difficulties in autism, where individuals may experience “mindblindness”, an inability to perceive others’ intentions, leading to profound social deficits(*151*, *152*). Using our network approach, we show that brain-wide networks in ASD-like mice exhibit reduced capacity to predict partner identity, indicating a partially impaired ToM in autism. Notably, to reach a firm conclusion about ASD, future studies should include additional mouse models carrying different mutations. Future work could extend this framework to compare brain-wide network dynamics during social interaction in models of other neuropsychiatric disorders, including schizophrenia, depression and post-trauma, and further quantifying the ability of different pharmacological (or non-pharmacological) treatments in restoring network deficits. In summary, our findings reveal coordinated brain-wide subnetworks that collectively encode critical aspects of social behavior across two interacting animals. These insights are pivotal to understanding the nature of social interaction and have far-reaching implications for both neuroscience and clinical research.

## Methods

### Animals

All experiments were approved by the Institutional Animal Care and Use Committee (IACUC) at the Hebrew University of Jerusalem, Israel (Permit Number: MD-20-16237-4). For experiments with wild-type mice (WT; Figures 1-4) a total of 23 C57BL/6 male mice (age 2–6 months) were used in this study. Experiments were done on groups of mice (3 groups of 5 and two groups of 4) that were housed together from the age of weaning (∼3-4 weeks). To study autism (Figures 5-6, we used 24 mutant SHANK3 mice (ASD mice; 4-8 weeks males; *Shank3*^Δ*4-22*^ mouse model; homozygous)(*31*, *153*) which is a well-established mouse line displaying several behavioral and social deficits (*36*, *48*, *53*, *154*, *155*). In short, SHANK3 is a postsynaptic scaffolding protein critical for the structure and function of excitatory synapses, especially in the postsynaptic density (PSD) of glutamatergic synapses, and mutations in SHANK3 are among the most strongly associated with ASD.

### Multi-fiber implant surgery

Mice were anesthetized with 2% isoflurane (in pure oxygen). Body temperature was maintained at 37 °C. To prevent inflammation and pain during anesthesia, we injected 0.1 μl/g body weight meloxicam subcutaneously. Connective tissue was removed from the skull, which was additionally polished and dried. To ensure optimal adhesion of the skull to the connective dental cement, we applied iBond (Kulzer, Total Etch). Small, slit-like craniotomies were made to allow for virus injections and implantation of fiber arrays (carried out on the same day). First, ∼200 nl of AAV.Syn.GCaMP6f.WPRE.SV40 were injected at 1.5 nl/sec through nanofil injector and nanofil needle (35BV inner diameter diameter: 55 μm) into 24 regions of interest. To allow local diffusion and avoid possible refluxes, we kept the needle in place for 2 min after injection. For the anterior array (12 fibers) was implanted at 1.54 mm anterior to bregma, and from 0.9 to 3.5 mm away from the midline, tilted at an angle of 20°. The posterior array (12 fibers) was implanted 1.7 mm posterior to bregma, and 1.8 to 4.4 mm away from the midline, tilted at an angle of 20°. Next, we applied dental cement (Tetric EvoFlow, T1) on the skull and around the implant followed by ultraviolet-light curing. A lightweight metal head post was cemented to the skull, allowing for head-fixation during the connection of the fiber bundle. After 2 weeks of recovery, mice were habituated for short head-fixation and moving freely within the cage while connected to the fiber bundle.

### Design of multi-fiber arrays

We utilized components of fiber connector technology adopted from Sych et al. 2019, specifically, ferrules that can accommodate 12 fibers (No. 17185, US Conec). Optical fibers with polyimide coating (100-μm core diameter, 124-μm outer diameter,0.22-NA, No. UM22-100, Thorlabs) were fitted into the ferrule. For targeting specific subsets of brain regions, fibers were precisely positioned at different length using a template printout of fiber. After alignment, fibers were glued to the ferrule, polished on the end side and alignment pins were added (No. 17799, US Conec).

### Multi-fiber photometry setup

Fiber bundle cables connect the ferrule in the mouse to the optical setup. One bundle reverted four ferrules of 12 fibers each (two ferrules from each mouse) into one 48 fiber (12 by 4) output going into the optical setup (Sylex s.r.o No. 1-830-830-480-380/00/0005.00). The optical setup is detailed in (*51*). In short, a Coherent OBIS LX 473-nm laser used for excitation is interleaved with a Coherent OBIS LX 405-nm that is used for control (capturing non-calcium related signal and movement artifacts; Frame rate of 20 Hz., 10 Hz. for each laser). A set of optics were used to create appropriate illumination patterns at the object plane of the objective. First, the beam was expanded and collimated (GBE05-A, Thorlabs). Then, a rectangle-like illumination pattern that fits the fiber array was created using two cylinder lenses (LJ1703RM-A and LJ1629RM-A, Thorlabs) placed before a dichroic cube (F38-495, AHF) and the objective (No. TL4x-SAP, Thorlabs). Power output was 9-20 µW from each ferrule (an average of 12µW per fiber). Excitation and emission filters were used to reduce residual broad spectrum light (474/27 No. F39-479SG for 473; 400/40 No. F39-400SG for 405; 525/50 nm, No. F37-516 for green emission, AHF). The fiber array end face was focused onto a camera sensor (ORCA Flash4.0 v3, Hamamatsu camera) with a tube lens with internal focusing (Proximity Series InfiniTube, 200-mm focal length).

### Behavior

Each group was composed of 4-6 cage mate mice (for WT: 3 groups with 5, 2 groups with 4; For ASD: 2 ASD groups, 5 mixed groups; Fig. S7. d). All mice with the group were implanted with a multi-fiber implant. Each mouse was identified by a different color on its head plate and dental cement. Within a recording day, all possible pairs (6-15 pairs per group, total 42 pairs for WT, 34 pairs for ASD) were recorded together and each group was imaged for 12±1 days (mean±SEM). Each pair was recorded for 10 minutes. In parallel to multi-fiber photometry a camera monitored from the top the position of both mice in the cage (30 Hz.; The imaging source; DFK 33UX273). To synchronize between the imaging and behavioural cameras, a green LED that was visible in the behavioural camera was triggered from the software of the imaging camera (LABVIEW 2019; NI) at the beginning and end of each 10-minute recording session.

### Data analysis

#### Social epoch definition

Mice were tracked offline using DeepLabCut software(*52*), with nine body-point labels per mouse, manually validated for switches in animal labels and further processed for outliers removal. The minimal distance (Euclidean) between two mice (out of the nine point) was calculated for each time frame and smoothed (moving window of 50 points; Fig. 1b). We defined a social interaction if the distance was under 8 cm or for at least 0.5 sec (also termed as ‘contact’ epochs; n=11,912 and 6722 for WT and ASD respectively). This threshold was chosen based on viewing social dynamics within the cage. ‘no-contact’ epochs (n=12,224 and 34729 for WT and ASD respectively) were defined when the distance was above 15 cm. In total the mice spent 126±36 (WT) and 201±77 (ASD) seconds per session on social contact (mean± SEM; Fig. 1d) and an average total of 2382 social interactions were detected for each WT group (ranging from 123 to 516 for each pair) and 955 for each ASD group (ranging from 59 to 392 for each pair).

#### Rank estimation

For each social interaction, we determined which mouse initiated the interaction (i.e., initiator) based on two parameters: (1) calculating which mouse travelled the greatest distance up to the interaction point (within 5 seconds preceding contact), and (2) using both mice body axis (i.e. if the nose of one mice was mostly close to the tail of the other during the beginning of the interaction). Visual inspection in a subset of videos validated the correctness of this definition. To determine the social rank of each mouse within its own group, we calculated the ratio for each pair between initiating a social interaction or being approached (i.e., approach ratio; Fig. 1e). The rank for each mouse was defined as the average approach ratio with all its partners (Fig. 1f). This social rank corresponded to other behavioral parameters that were tested such as: chase behavior, velocity, cage occupancy, elevated plus maze test and open field test (Fig. S2).

##### *Chase Behavior* (Fig. S1. a)

For each social interaction, a chase was defined under the following criteria: 1. The nose-to-tail distance (between pairs) during the first two-thirds of the interaction was the shortest compared to other distances (e.g., nose-to-body or nose-to-nose). 2. The velocity of both mice was greater than 10 cm/s. 3. The total distance traveled during the interaction exceeded 10 cm. Dominant mice are predicted to engage in more chase behaviors.

##### *Cage occupancy* (Fig. S1. c)

We divided the behavioral cage into 1 cm² grid squares. Then, for each square, we determined whether the mouse passed through it or not. Based on this, we determined the total number of square centimeters the mouse occupied divided by the total cage area. Dominant mice are predicted to have a higher cage occupancy.

##### *Open Field* (Fig. S1. d)

Groups #3 and #5 in WT groups and groups 1, 4-7 in ASD groups underwent the open field test (n=7 tests/mouse). During each test, mice freely explored a closed 50x50 cm arena for 6 minutes. The edges of the arena were defined as the 7 cm perimeter, while the remaining central area was designated as the open area. Mice tend to spend more time at the edges of the arena, especially anxious mice. Thus, a high open/edge ratio (in terms of time spent in each part) indicates a less anxious mouse which is predicted to have a higher social rank.

##### Elevated plus maze (Fig. S1. e)

Groups #2 and #3 in the WT groups and all ASD groups underwent an elevated plus maze which is an anxiety-related test), comprised of an elevated platform with four arms, two closed and two open (n=7 for each mouse; 15 minutes each. Mice tend to spend more time in the closed arms, especially anxious mice. Thus, a high open/close ratio (in terms of time spent in each part) indicates a less anxious mouse which is predicted to have a higher social rank.

#### High and equal rank ratio pairs

We further divided all pairs (n=42) into a high rank ratio group (n=20 approach ratio<1.35, ranging from 1.39 to 2.73) and an equal rank ratio group (n=20 approach; ratio>1.35, ranging from 1 to 1.349; Fig. 4a) in the WT groups. In the ASD groups the high rank ratio group (n=19 approach ratio>1.5 ranging from 2.8 to 1.51) and the equal rank ratio group (n=15 approach; ratio<1.5, ranging from 1 to 1.45). The rank ratio was determined based on the approach/being approached ratio between each pair. Within each group, we focused on social interactions in which the higher ranked mouse initiated a social interaction and approached the lower ranked mouse (WT groups: dominant: 170 ± 59 interactions, subordinate: 101 ± 36 ASD groups: dominant: 106±42 subordinate 94±46 epochs for the high rank ratio pairs; WT groups: Dominant: 146 ± 51, subordinate: 124 ± 42 for the equal rank ratio pairs. ASD groups: dominant: 117±28 subordinate: 104± 32).

#### Multi-fiber pre-processing

Data analysis was performed using Matlab software (Mathworks). Multi-fiber fluorescence images (48 fiber tips at the end plate of the ferrule) were converted to Matlab files and pixels within each fiber tip were averaged and labelled as a specific brain area. Each fiber signal was smoothed (moving average window of 5 frames) and divided by its median across the session (10 min; ΔF/F). Next, artefacts (defined as signals beyond 35% ΔF/F) were removed and each signal was linearly detrended. In recording sessions where a control light (405 nm) was interleaved with the 473 nm light (sampled at 20 Hz, 10 Hz per channel), we calculated the ΔF/F for each channel separately and then subtracted the 405 control signal from the 473 signal. Next, brain signals were synchronized to the behavioural video using a green LED that was triggered by the fiber camera and captured in the behavioural videos. Examples of post-processed and synchronized brain signals are presented in Figure 1j. Fiber signals can further be aligned to social contact and averaged across social epochs within each mouse and across mice (Fig. 1k)

#### Classifying social interaction

For each possible pair, we divided fiber signals in 24 brain areas into social interaction epochs (i.e., ‘contact’) and non-social epochs (i.e., ‘no-contact’). A support vector machine (SVM) with a polynomial kernel classified ‘contact’ or ‘no-contact’ epochs based on signals in 24 brain areas. The classifier was trained on 80% of the data (259±8 for ‘contact’ and 1312±52 for ‘no contact’) and the remaining 20% were used as a test. The training and testing data were stratified to account for the imbalance between the two groups, and each fiber’s signal was normalized before training. Classifier performance was assessed by the area under the curve (AUC) of a receiver operating curve (ROC) derived from the test data (Fig. 1n). The procedure was cross validated for each mouse (n=10). As an additional control, we performed a similar analysis on trial shuffled data (n=100 iterations).

#### Neuronal Controls for social interactions

##### Unpaired control

To control possible common input during social interaction, we performed the network correlation analysis on randomly shuffled pairs of mice (i.e., unpaired control; n=42). Each epoch in one mouse was matched with a random epoch in a different mouse, while maintaining the temporal statistics. This was done for between and within brain correlations and the differential correlation matrix was calculated and clustered (Fig. 2b and Fig. S5. e-h).

##### Movement control

To control whether enhanced responses during social contact are explained by increased movement of the mice, we divided ‘non-contact’ based on the velocity of the mouse, into slow (velocity < 1 cm/sec) and fast (velocity > 4 cm/sec) epochs. This was done for each mouse (n=23) and response difference between ‘contact’ and between ‘no-contact’ fast and ‘no-contact’ slow were calculated (Fig. S3. a-d). In the former case results were similar compared to taking the whole ‘no-contact’ period whereas in the latter case there were minor differences between fast and slow epochs.

##### Non-social object control

To control weather brain responses could arise from sensory integration of a non-social stimulus, 16 mice were put into an arena with two objects and were allowed to freely explore. “Object contact” was defined as epochs that the mouse was within 4.5 cm distance from the object (20±7% of time; mean±std), and “object no contact” were defined as the remaining time. A similar subtraction in response between ‘contact’ and ‘no-contact’ epochs was performed resulting in minor and insignificant differences across the brain (Fig. S3. e).

##### Separate cages control

To control whether similar results can be obtained without social contact, nine pairs of mice were simultaneously imaged while exploring adjacent separate cages divided by a high and sealed partition. We divided the data into pseudo-’contact’ epochs in which the mice were close in distance (<20 cm, n=27±3 epochs, 4.7±1.7 seconds duration, mean±SEM) and pseudo-’no contact’ epoch in which the mice were far from each other (>25 cm, n=45±3 epochs, 6.3±2.4 second duration). Next, the difference in activation was calculated for each area resulting in minor and insignificant differences across the brain (Fig. S3. f)

##### Random ‘no contact control’

To control non-socially related correlations, we randomly split the “no contact” epochs into two groups and computed the correlation difference matrix (Fig. S5e-g). This was done for each pair (n=42) and the differential correlation matrix was calculated, sorted and clustered, resulting in non-meaningful clusters (Fig. S5. h).

##### Network correlation analysis

To investigate brain-wide interactions within and between and within brains, we performed a correlational network analysis. For each recording session of a given pair, we calculated the full correlation matrix (Pearson’s correlation coefficient) between all possible pairs of brain areas (48 by 48 from both brains) for each ‘contact’ (n=11,912 length= 4.56±10.92 sec at least 1 second duration) and ‘no-contact’ epoch (n=12,056 length=9.125±11.58; at least 1 second duration). Full correlation matrices were then pooled across epochs (median) within each mouse (n=42; Fig. S4. a). The same analysis was done for the ASD pairs (n=6722 contacts duration= 4.65±39.7 seconds n=7611 ‘no-contact’ duration= 9±44.9 seconds; median±std). For each pair, we kept the higher ranked mouse on the top right quadrant and the lower ranked mouse on the lower left quadrant. Next, we calculated the differential correlation matrix by subtracting the correlation matrix during ‘no-contact’ from the ‘contact’ matrix (i.e., Δr; Fig. 2a). Positive Δr values indicate a bias towards social contact whereas negative values indicate a bias towards periods outside social contact. We note that in this type of analysis an increase in correlation values, even in negative values (e.g., an increase from −0.6 to −0.4 will result in a positive Δr) is considered as bias toward social contact. The logic behind this is that we assume that negative values indicate that two brain areas belong to separate subnetworks and therefore should be clustered separately. Finally, differential correlation matrices were averaged across mice pairs (n=42 for WT pairs, n=34 for ASD pairs; Fig. 2a, Fig. 4e)

Next, we wanted to cluster the differential correlation matrix (averaged across mice) into distinct subnetworks. Next, we used a hierarchical clustering algorithm (Ward’s method, Euclidean distance) and plotted the cluster tree as a dendrogram (Fig. 2c). Based on the distance between branches, we chose to cluster the matrix into 3 clusters. We note that there may be additional subnetworks within each cluster that can encode unique social information, for example two distinct clusters within cluster 3 or 2. Next, the strongest 30% Δr for each cluster can be superimposed back on the brain slices where each line is color coded for the sign of the Δr and its width indicates its magnitude (Fig. 2f). A similar presentation can be done for between clusters Δr values (Fig. S4. d; strongest 20%). To assess the stability of our clustering algorithm we used a bootstrapping method, in which we performed a similar clustering pipeline for only 80% of the data over numerous iterations (n=1000). For each brain area we compared the identity of its cluster for each iteration with the observed cluster identify and calculated how many iterations had a correct match (%; Fig. 2i). If a certain brain area always belonged to the same cluster, then it was considered stable. The identity of each brain area to each of the 3 clusters is plotted in Fig. S4. f. Most brain areas display stable cluster identity (>80%) whereas few areas were unstable. For example, the BLA of the dominant mouse may be better identified with cluster 1 instead of cluster 2. Finally, similar analysis using t-SNE products gave similar brain clusters (Fig. S4. b,c).

##### Asymmetry index between dominant and subordinate mice

To investigate brain-wide network differences based on social rank, we defined a social asymmetry index (ASI). For each pair of brain areas (i.e., connection) the ASI was defined as the difference in Δr between the dominant and subordinate mice divided by the sum of their absolute values (Fig. 3a). The ASI was then averaged across all 42 pairs (Fig. 3b). Positive ASI values indicate a bias for the dominant mouse whereas negative ASI values indicate a bias towards the subordinate mouse. Furthermore, ASI values can be divided into within and between connections and also into the clusters from the network analysis.

##### Hub brain areas

To identify potential hub regions within brain-wide subnetworks, we extracted for each brain area its maximal Δr value. This was done separately for within and between brain connections. Thus, each brain area is left with one maximal connection within its own brain and between the partner’s brain. For example, Figure 3e displays the maximal Δr value for brain areas in the dominant mouse. From here, we can calculate the number of maximal connections received in each brain area of the subordinate mouse. This was done for each pair and averaged across pairs (Fig. 3f-h). A similar approach was done for within brain maximal connections (Fig. 3i-l). Thus, brain areas that receive many maximal connections are considered as hub areas.

##### Non-matching index

To quantify the relationship between a certain brain area and its connection with homologous (matching) or non-homologous (non-matching) brain areas in the partner brain, we defined a non-matching index (NMI; Fig. 3m). The NMI was defined as the difference between the maximal non-homologous Δ*r* and homologous Δ*r*, divided by the absolute sum of the two. The NMI was calculated for each pair and averaged across all 42 pairs (Fig. 3p). Positive NMI values indicate a bias towards non-homologous interactions whereas negative NMI values indicate a bias towards homologous interactions. We note that NMI is inherently biased towards non-homologous interactions (since it takes the maximal non-homologous value out of 23 possible connections), but in our case, we wanted to make a general point that interactions between non-homologous brain areas are of high interest and display strong inter-brain connections (as compared to homologous connections).

##### Classifying initiator/being approached

An SVM with a gaussian kernel was trained to classify whether a mouse initiated a social interaction or was being approached, based on brain signals from each mouse (Fig. S6. a,b). Here we focused on a dominant mouse approaching the subordinate mouse. This was done separately for high and equal rank ratio groups (n=20 and 22 mice for high and equal rank groups respectively; n=3446 and 2984 social interactions for high and equal rank groups respectively) and within three distinct time periods in relation to social contact: before (−2.75 to −1.5 s), during (−1 to 1 s), and after (2 to 6 s). Th classifier was trained on 80% of the data and tested on the remaining 20%. The procedure was cross validated (n=10) and we performed a similar analysis on trial shuffled data (n=100 iterations).

##### Classifying mouse identity

An SVM with a gaussian kernel was trained to classify the identity of the partner mouse based on the brain of one mouse (Fig. 4e). To do this, the brain signal during social contact for each mouse (n=23) was divided based on the interacting partner. The classifier was trained (80%) to identify which mouse was in contact for each social interaction (872 ± 326 (mean ± SD) social interactions from 3 or 4 partners for each mouse) and corrected for unequal group size. Accuracy was tested on the remaining 20%. The procedure was cross validated (n=10) and we performed a similar analysis on trial shuffled data (n=100 iterations). For each mouse, a confusion matrix can be plotted to detail the accuracy of the classifier for each partner (Fig. 4h). The confusion matrix is then averaged across mice by averaging the relative accuracy (compared to its trial-shuffled accuracy) across mice for each cell in the confusion matrix (Fig. 4i).

### Histology

Mice were euthanized with an overdose of isoflurane and were perfused transcardially with phosphate-buffered saline (PBS) followed by 4% paraformaldehyde (PFA) in PBS. After perfusion, tissue was removed from the skull, and the head, including the multi-fiber implant, was additionally fixated in 4% paraformaldehyde for 1 week. The ventral (bottom) side of the skull bone was removed and the brain was carefully extruded. Coronal sections (80–90 μm) were cut with a vibratome (No. VT100, Leica), mounted onto glass slides, adhered and colored with DAPI (4′,6-diamidino-2-phenylindole) and imaged using Nikon SMZ25 fluorescent microscope (Fig. 1h). n most mice, several fiber tips were clearly identifiable. Using these, along with the known coordinates of the fiber array, we estimated the positions of the undetected fibers. In total, we successfully localized 24 fiber tips in 19 out of 23 mice, as shown in Fig. S2. The average localization accuracy was 80%.

### Statistical analysis

In general, the Wilcoxon signed-rank test was used to compare a population’s median to zero (or between two paired populations). For non-paired populations we used a Mann– Whitney U-test to compare between medians. Multiple group correction was used when comparing between more than two groups. In all decoding analysis (e.g., SVM) a cross validation and a trial shuffled control was performed and compared to the observed data.

## Data availability

Data will be shared upon reasonable request by the corresponding author. No new material was generated in this work.

## Author contribution

O.M. and A.G. conceptualized the study. O.M. and A.G. performed the experiments, preprocessed and analyzed the data. R.T and V.K. contributed to surgeries, histology and other assistance. O.M., H.A. and A.G. wrote the manuscript, S.G. handled the ASD mice and all authors revised, read and approved the final version of the manuscript.

## Acknowledgements.

We would like to thank Dr. Yaroslav Sych for helping with the multi-fiber setup, and Eden Rosenstein for implementing the multi-SVM code for identity detection. We thank Bar Peled for manually validating identity switches in animal labels within the DeepLabCut-generated trajectories. We thank Doron Michaeli for helping with the ASD line and running behavioral tests. This work is funded by the US Department of Defense through the Congressionally Directed Medical Research Programs (CDMRP; AR210106; A.G.) and the European Union (ERC Starting Grant, MESO-AG, 101040378; A.G.).

