## Supplemental Figures and legends for "Brain-Wide Subnetworks within and between Naturally Socializing Typical and Autism Model Mice"

Supplementary Figures 1-9:

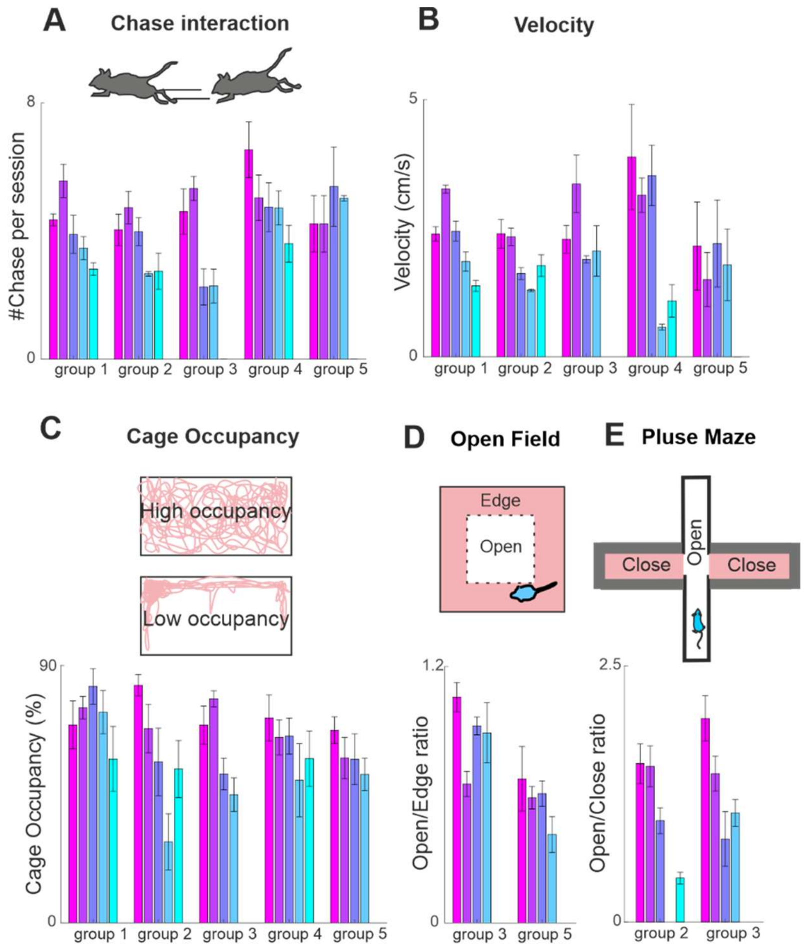


**Fig. S1.**

**Validations for social rank. a.** Chace behavior: average number of chases per session, for each mouse ordered by the social rank defined in Figure 1f. Error bars depict SEM across pairs. **b.** Velocity: average velocity within the behavioral cage for each mouse ordered by the social rank defined in Figure 1f. Error bars depict mean±SEM across pairs. **c.** Cage occupancy: mean occupancy (percentage of area covered) within the behavioral cage ordered by the social rank defined in Figure 1f. Error bars depict mean±SEM across pairs. **d.** Open field test: Several mice underwent an open field test and the average time spent in the open vs edges (ratio) were calculated for each mouse and ordered by the social rank defined in Figure 1f. Error bars depict mean±SEM across sessions (n=7). **e.** Elevated plus maze: Several mice underwent an elevated plus maze and average time spent in the open vs closed arms (ratio) was calculated for each mouse and by the social rank defined in Figure 1f. Error bars depict mean±SEM across sessions (n=7).

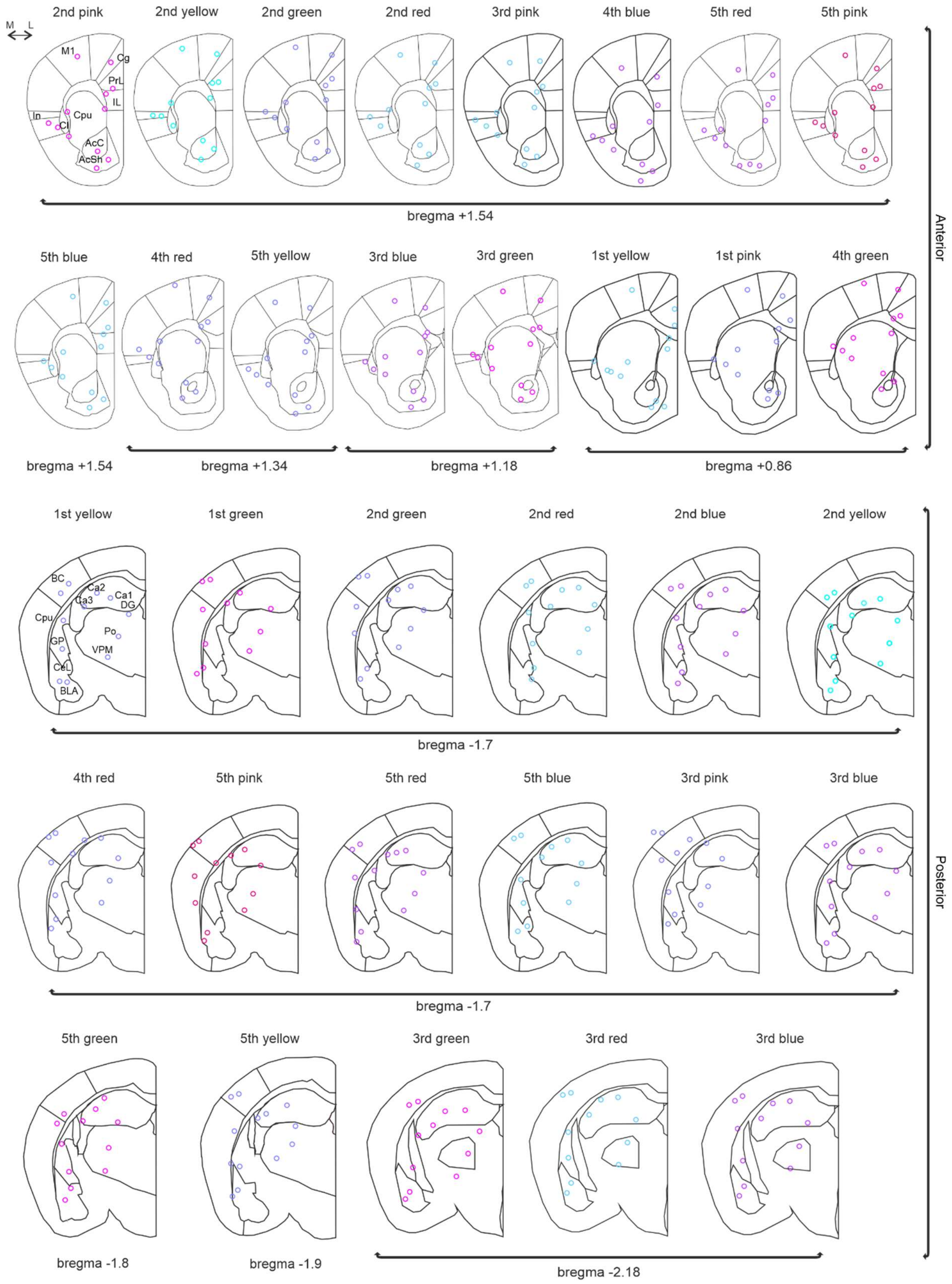


**Fig. S2.**

Fibers tip locations based on histology for 19 out of the 23 mice. Colors indicate social rank.

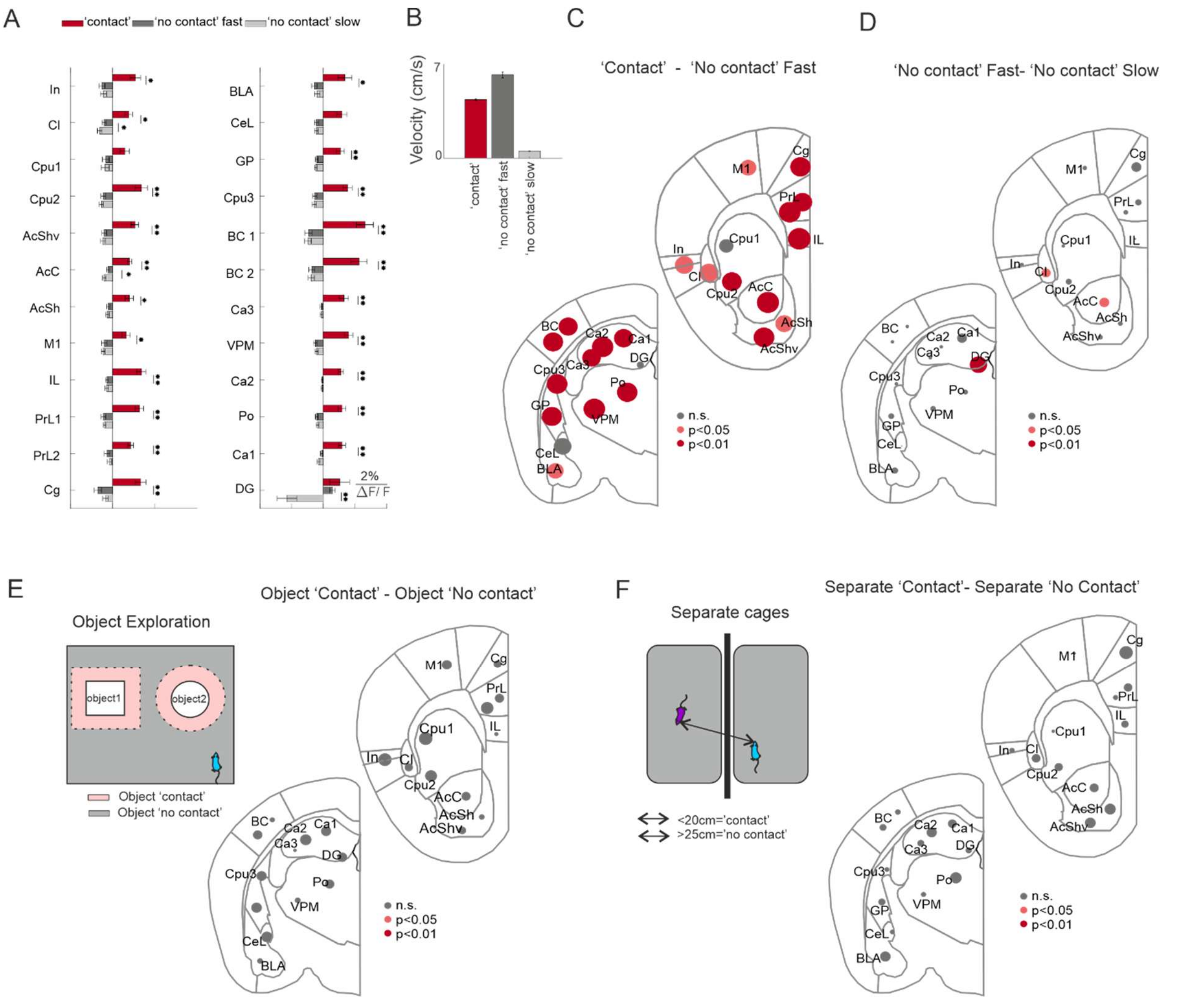


**Fig. S3.**

**Controls for brain-wide social responses: a-c Movement control. a.** 'No-contact' epochs were divided into fast and slow based on the velocity of both mice (see Methods). Average responses (ΔF/F) during social interaction ('contact'; red), 'no-contact fast' (dark gray) and 'no-contact slow' (light gray) are plotted for each brain area. Error bras depict mean±SEM across mice (n=23).**$p<0.01$,*$p<0.05$, Wilcoxon signed-rank test with Bonferroni correction. **b.** Average velocity during 'contact' (red), 'no contact fast' (dark gray), and 'no contact slow' (light gray). Error bars as in a. **c.** Circle representation depicting the difference in responses between 'contact' and 'no contact fast' periods divided by the sum SEM of both (as in Fig. 2d). The larger the circle the bigger the distance between contact and no-contact periods. Significance levels are color coded. **d.** Same as c but for the difference between 'no-contact fast' and 'no contact slow'. **e. Object contact control.** Mice were put in a behavioral cage with two object and were allowed to freely explore. Object 'contact' and 'no-contact' epochs were defined (see Methods). Circle representation depicting the difference in responses between 'object contact' and 'object no-contact'. **f.** Separate cages control. Nine pairs of mice were imaged while freely moving in separate adjacent cages and 'pseudo-contact' and 'pseudo no-contact' epochs were defined based on their distance (see Methods). Circle representation depicting the difference in responses between 'pseudo contact' and 'pseudo no-contact'.

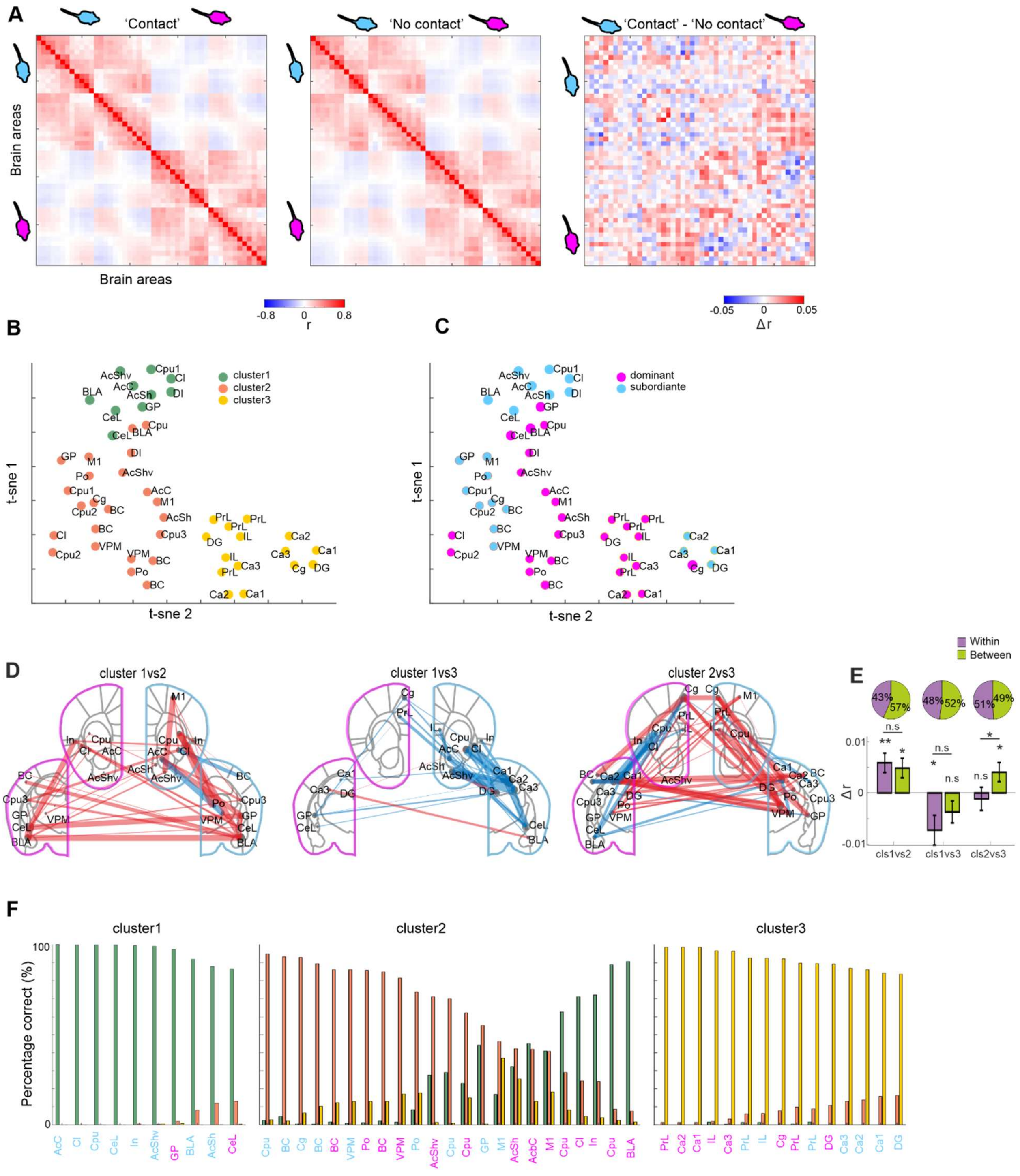


**Fig S4.**

**Additional network and clustering data. a.** Full correlation matrix for 'contact' (left) and 'no-contact' (right) epochs separately, averaged across mice pairs (n=42) and sorted for dominant and subordinate mice. Differential matrix as in figure 3a is presented on the left. **b**. As an alternative clustering method, 2D embedding (t-SNE) of the differential correlation matrix (Fig. 3a; correlation as a distance measure). Color coded for the three clusters obtained by the original clustering method from Fig. 3c. **c.** Same as in b but color coded for dominant and subordinate mice

**d.** Cluster assignment stability for each fiber, quantified by bootstrap resampling (see Methods) and plotted for each of the 3 clusters. High values depict high stability in cluster assignment.

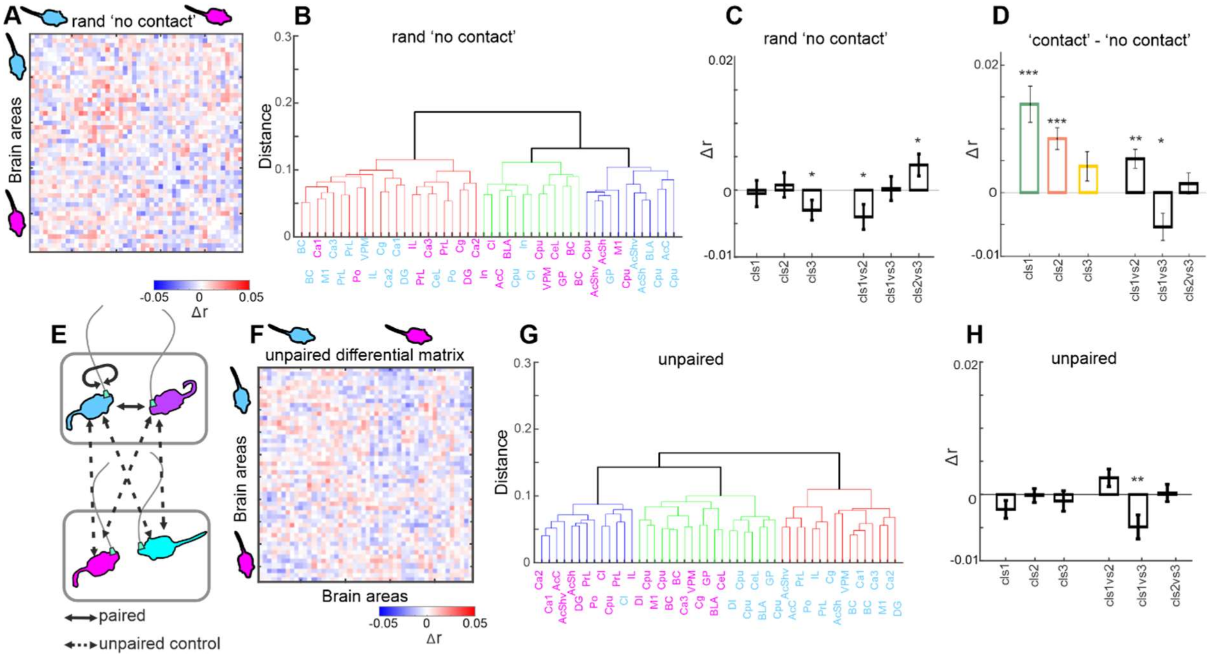


**Fig S5.**

**Controls for network analysis. a-d.** Random epoch correlation control: **a.** Differential correlation matrix obtained by subtracting correlation matrices from two randomly selected groups of 'no-contact' epochs (see Methods). **b.** Hierarchical clustering dendrogram based on the matrix in e. Three clusters are color coded. **c.** Δr values in each cluster and between clusters. Error bars depict mean+-SEM across pairs (n=42). **d.** Observed Δr values as in figure 2e displayed for comparison. **e-h.** Unpaired correlation control. **e.** Schematic illustration of the "unpaired control" (see Methods). **f.** Average differential correlation matrix derived from the "unpaired control". **g.** Dendrogram from hierarchical clustering on the matrix in j. **h.** Δr values in each cluster and between clusters in the unpaired control. Error bars depict mean+-SEM across pairs (n=42).

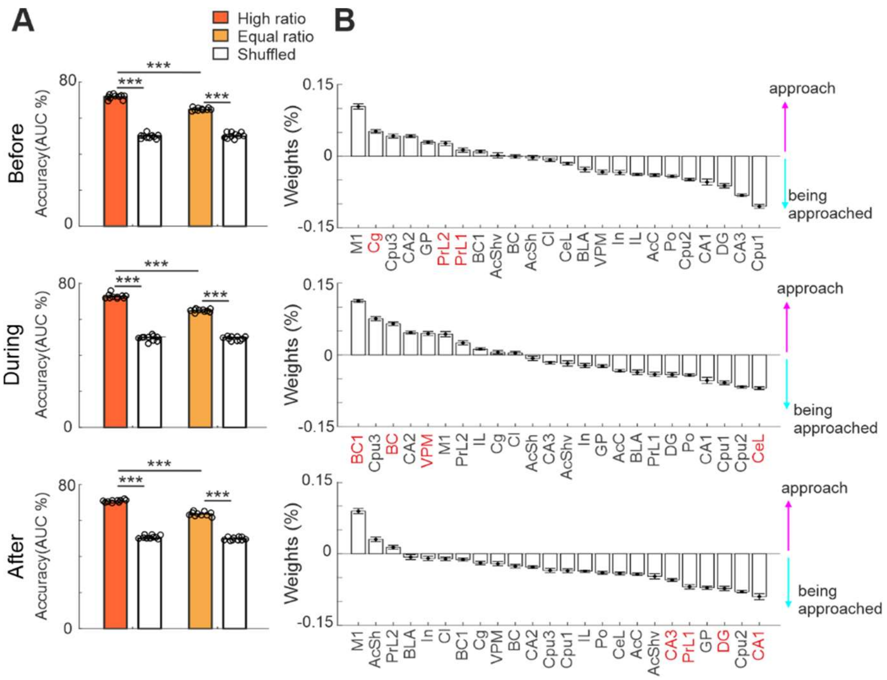


**Fig S6.**

**Predicting initiation of social contact based on brain-wide dynamics. a.** Accuracy of an SVM classifier predicting whether a mouse initiated a social interaction based on different time periods: before (top), during (middle) and after (bottom) social contact (see Methods). This was done for high and equal rank pairs separately. Accuracy of the SVM on trial shuffled data is presented in empty bars. dots depict each 10-fold run. **f.** The weights (sorted across brain areas) of the corresponding classifier in a. Error bar depicts mean±SEM across 10 fold repetitions. Areas of interest are colored in red.

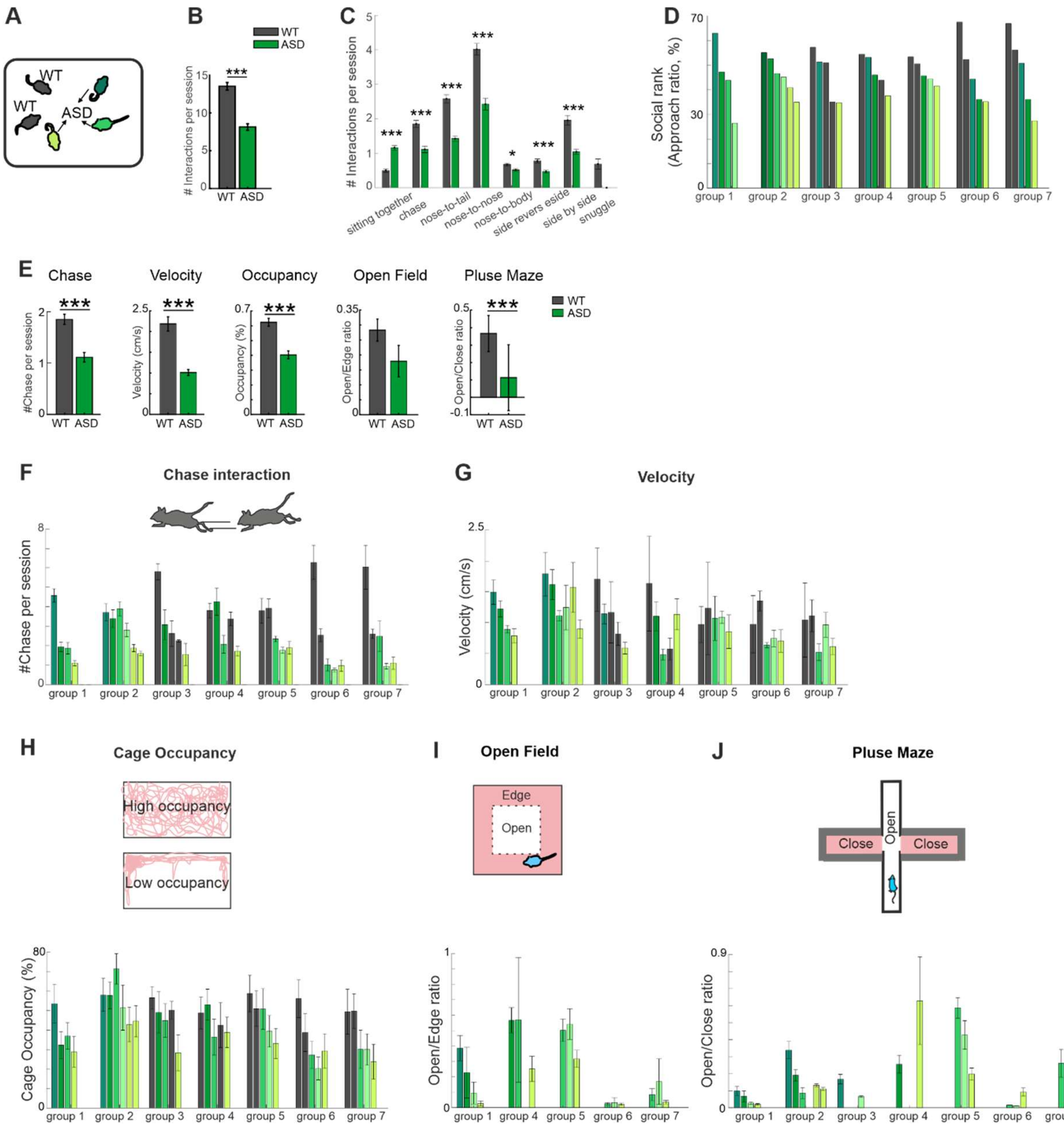


**Fig S7.**

**Validations for ASD behaviors**. **a.** Schematic illustration of behavioral paradigm for ASD mice. **b.** Number of initiating social interactions for ASD (green) and WT (gray) mice. error bars depict mean±SEM across mice (n=98 and 68 for ASD and WT mice pairs respectively). **c.** Number of social interactions for ASD (green) and WT (gray) mice. Error bars as in b. **d.** Relative rank of each group (groups 1 and 2 only ASD mice; Groups 3-7 mixed with WT) based on the initiation ratio (as in Fig. 1d). Colors denote relative rank. **e.** Wilde type and ASD like mice comparison of several behavior parameters**.** Error bars depict mean±SEM across mice (chase, velocity, occupancy n=23 and 24 for Wilde type and autistic mice respectively. Open field n=17 and 16, plus maze, n= 31 and 24. **f.** Chace behavior: average number of chases per session, for each mouse ordered by the social rank defined in Figure 1f and here in d. Error bars depict SEM across pairs. **g.** Velocity: average velocity within the behavioral cage for each mouse ordered by the social rank defined in Figure 1f. Error bars depict mean±SEM across pairs. **g.** Cage occupancy: mean occupancy (percentage of area covered) within the behavioral cage ordered by the social rank defined in d. Error bars depict mean±SEM across pairs. **h.** Open field test: the autistic mice underwent an open field test and the average time spent in the open vs edges (ratio) were calculated for each mouse and ordered by the social rank defined in d.  Error bars depict mean±SEM across sessions (n=7). **e**. Elevated plus maze: Autistic mice underwent an elevated plus maze and average time spent in the open vs closed arms (ratio) was calculated for each mouse and by the social rank defined in d. Error bars depict mean±SEM across sessions (n=7).

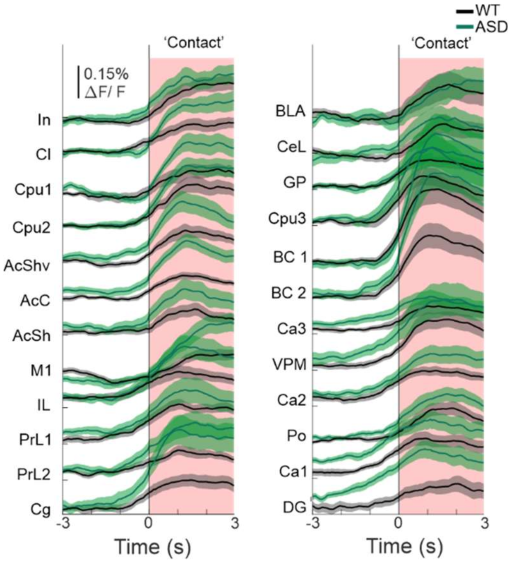


**Fig S8.**

Responses (ΔF/F) in 24 brain areas aligned on social contact (time 0) for WT (black) and ASD (green) mice. Error bars depict mean±SEM across mice (n=23 and 24, respectively).

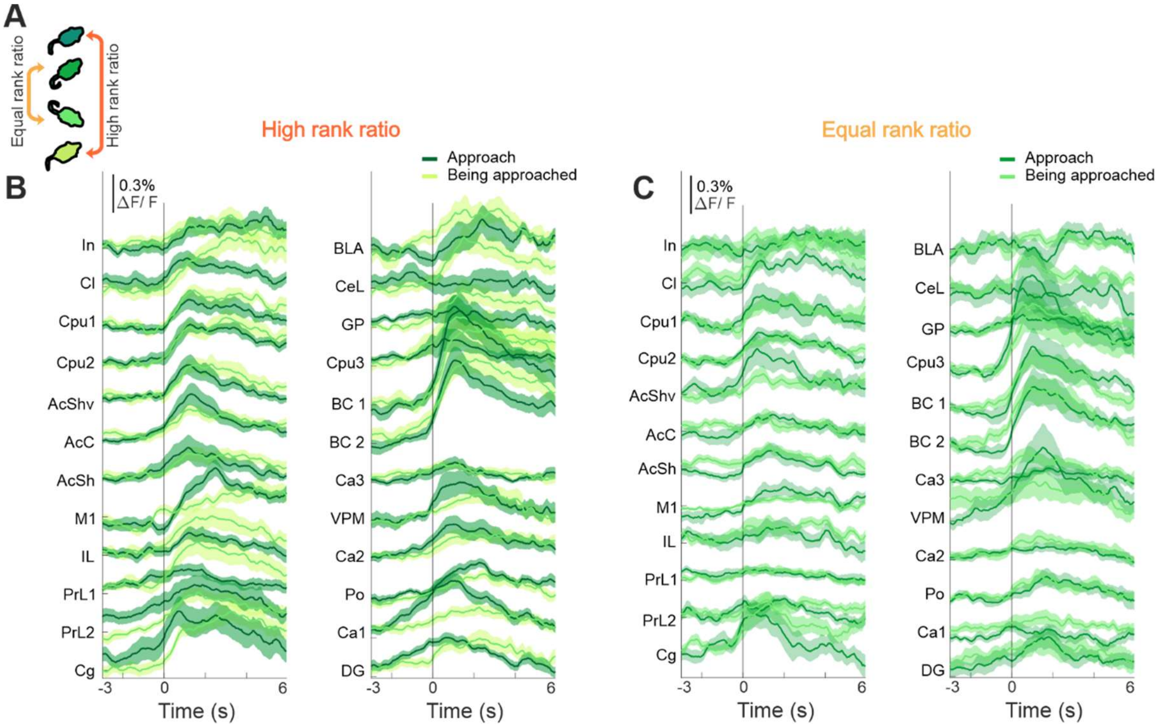


**Fig S9.**

**Average traces in 24 brain areas divided to dominant approach/subordinate being approached in high ratio and equal ratio groups**. **a.** Schematic of high and equal rank ratio pairs and focusing on interaction upon which the higher ranked mouse initiated social interaction. **b**. Neuronal responses in 24 brain areas (divided into clusters) for high rank pairs. Error bars depict mean±SEM across pairs (n=19). **c.** same as b. for equal rank pairs (n=15).
